# Mouse oocytes develop in cysts with the help of nurse cells

**DOI:** 10.1101/2021.11.04.467284

**Authors:** Wanbao Niu, Allan C. Spradling

## Abstract

Mammalian oocytes develop initially in cysts containing many more germ cells than the primordial oocytes they generate. We identified abundant nurse cells with reduced unique molecular identifiers (UMI)/cell from ovaries aged E14.5 to P1. Low UMI nurse cells are found in cysts and express the same major meiotic genes as pro-oocytes of the same stage, suggesting they are oocyte sisters that are signaled to transfer cytoplasm at different times and only subsequently diverge. Oocyte vs nurse cell selection occurs in cysts with a robust microtubule cytoskeleton, that closely interact with somatic cells and that develop a dense actin cytoskeleton around nurse cell nuclei that are held back from cytoplasmic transfer. Mouse and Drosophila nurse cells undergo programmed cell death by acidification from adjacent somatic pre-granulosa cells that express V-ATPases and cathepsin proteins. Disrupting acidification in cultured mouse ovaries blocked nurse cell turnover. About 200 genes are induced in mouse dictyate oocytes as previously reported, including Tuba1c and Tubb2b, genes that we find contribute to Balbiani body formation. Thus, mouse oocytes are specified within germline cysts and develop with the assistance of nurse cells using highly conserved mechanisms.

## Introduction

Mammalian female germ cells begin developing immediately after reaching the embryonic ovary, rapidly undergo key meiotic events and form into ovarian follicles by just five days after birth (Edson et al. 2009; Richards and Pangas, 2010; Lin and Capel, 2015; Ikami et al. 2017; Wu and Dean 2020). Pre-follicular development takes place in germ cell groups interconnected by intercellular bridges (IBs) known as cysts, stereotyped structures initiating both male and female gametogenesis prior to meiosis in diverse animals (King, 1970; Gondos, 1993; Pepling and Spradling, 1998; Matova and Cooley, 2001; Greenbaum et al. 2011; Haglund et al. 2011; Lu et al. 2017). Subsequently, most cyst germ cells turn over, while survivors develop a Balbiani body (Bb) (Cox and Spradling, 2003; Pepling et al. 2007) and become primordial oocytes (Pepling and Spradling, 2001; Lei and Spradling, 2013; 2016). Somatic pre-granulosa cells that interact closely with developing cysts and follicles also show evidence of evolutionary conservation (Stevant et al. 2019; Niu and Spradling, 2020). However, the importance of cysts to mammalian oocyte development and the role of the cyst cells that do not become oocytes has remained less clear.

Cysts play two roles in Drosophila and other animals where cyst function has been extensively investigated using developmental genetic and comparative studies (King, 1970; Spradling, 1993; Büning, 1994; Huynh and St. Johnston, 2004; Świątek and Urbisz, 2017). First, cyst production establishes an asymmetric microtubule cytoskeletal polarity based on its formative divisions (Koch and King, 1966; Lin et al., 1994; deCuevas and Spradling, 1998; Roper and Brown, 2004; Huynh and St.Johnston, 2004). This polarity underlies specification of distinct nurse cell and oocyte fates, and likely blueprints the earliest polarity of the oocyte itself. In Drosophila, the initial cyst cell will normally become the oocyte, while the other fifteen cells are selected to serve as nurse cells (Theurkauf et al. 1993; McGrail and Hays, 1997; deCuevas and Spradling, 1998). Shortly after the fourth mitotic cyst division, all 16 cells enter meiotic S phase. Subsequently, over several days, centrioles, mitochondria, Golgi, ER, cytoplasm and specific mRNAs move through the cyst’s nurse cells until they reach the oocyte (Mahowald and Strassheim, 1970; Grieder et al. 2000; Bolivar et al. 2001; Cox and Spradling, 2003). Organelles eventually enter the oocyte en masse and generate a Balbiani body, whose centrioles and some mRNAs relocate to define the oocyte posterior as the new follicle forms. Later, these nurse cells dump their remaining cytoplasmic contents into the oocyte, and undergo a distinctive pathway of programmed cell death (reviewed by Lebo and McCall, 2021).

Considerable evidence already argues that a fraction of mammalian cyst cells also act like nurse cells by transferring their cytoplasm and organelles into pro-oocytes. In lineage-marked mouse cysts, most of the centrioles, Golgi, ER, mitochondria and cytoplasm are transferred during meiotic prophase from cells fated to turn over and into surviving oocytes, aided by the partial breakdown of germ cell membranes (Figure 1A,1B; Lei and Spradling, 2016). In contrast, studies of the intercellular bridge protein Tex14 had showed that normal IBs are essential in developing male but not female gametes, calling the importance of cysts and oocyte nursing into question (Greenbaum et al. 2006; 2009). Recently, detailed studies confirmed that *Tex14* mutation disrupts normal IB formation, but showed that some female cysts still form and polarize (Ikami et al. 2021). Cyst cells compensate for the defective bridges and are still able to transfer organelles from nurse cells to oocytes through broken membranes, producing a reduced number of fertile oocytes. Moreover, disrupting IBs accelerated the kinetics of organelle transfer, suggested that IBs mediate a repressive signal (Ikami et al. 2021). However, the generality of nurse cell function has also remained uncertain. While cysts and Balbiani bodies are found in many vertebrates (Hertig and Adams, 1967; Heasman et al. 1984; Nakamura et al. 2010; Elkouby and Mullins, 2017) whether most cyst cells turn over and whether they contribute to Balbiani body formation has not been established (Kloc et al. 2004; Jamieson-Lucy and Mullins, 2019). Moreover, it has remained difficult to identify and study cyst germ cells that do not become oocytes except in insects.

**Figure 1.**
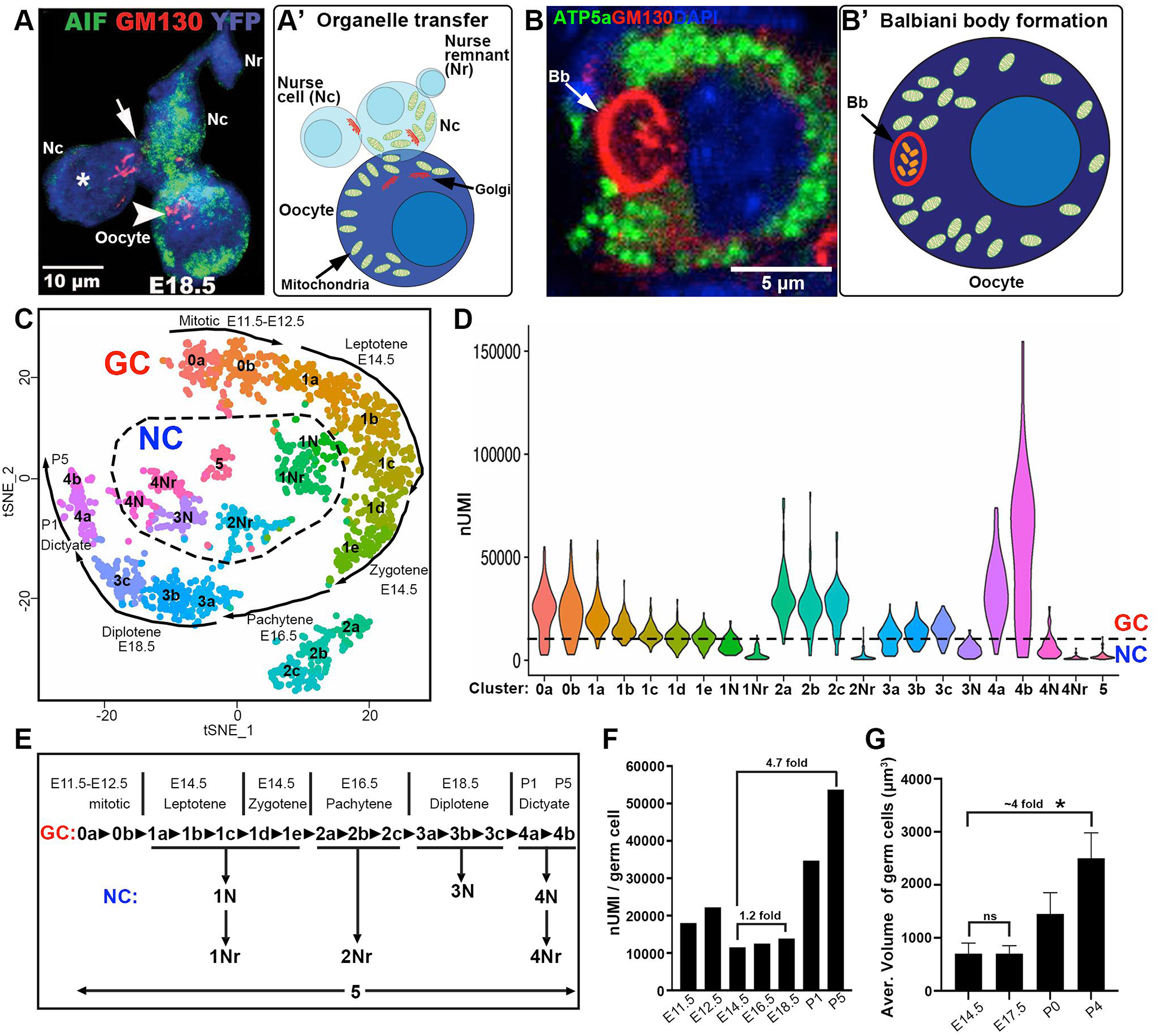
Identification of female germ cell scRNAseq clusters comprising oocytes and nurse cells. A. An E18.5 germline cyst lineage-labeled with YFP (blue) shows a tiny nurse cell remnant (Nr), and two other nurse cells (Nc) transferring mitochondria (AIF, green) and Golgi (GM130, red); into an oocyte (arrowhead). Image reproduced from Lei and Spradling (2016). A’. Diagram of the cells in A. B. A P4 oocyte showing a Balbiani body (Bb, arrow) with typical Golgi elements (GM130, red), and clustered mitochondria (ATP5a, green). (DAPI, blue). B’. Balbiani body diagram, showing its core of centrosomes (orange ovals), the Golgi shell (red), dispersed mitochondria (green) and nucleus (light blue). C. tSNE diagram generated of 22 germ cell (GC) clusters from mouse ovarian germ cells spanning E11.5-P5. Development from mitotic germ cells (0a) to dictyate oocytes (4a,b) proceeds around the outside as shown (arrows). Clusters of nurse cells (NC) are located within the dashed central area. The meiotic stage and developmental time germ cells are observed in the clusters are indicated near the arrows. D. Plot showing the larger UMI distributions of meiotic germ cells (clusters mostly above dashed line) compared to nurse cells (clusters mostly below dashed line). E. Summary diagram relating GC clusters (0-4b) to time points and inferred meiotic stages (above). Below, nurse cell clusters are positioned on the diagram based on their time detection and location in the tSNE plot. F. The total germ cell UMI in non-nurse cells at the indicated developmental ages up to P5. Fold differences are indicated. G. The average volume of germ cells as measured by Lei and Spradling (2016) at the indicated stages is shown for comparison to F. Size bars are indicated.

Here we report studies that clarify the central role of germline cysts in mammalian oocyte production and identify most cyst cells as nurse cells. Nurse cell transcriptomes determined using scRNAseq closely resemble those of pro-oocytes at the same stage, suggesting that mouse nurse cells do not differentiate into somatic cells, like Drosophila nurse cells do after follicle formation (Deluca and Spradling, 2020). The mouse cyst develops an extensive microtubule and actin cytoskeleton that we propose helps specify nurse cell vs oocyte fates in conjunction with internal and somatic signals. Moreover, mouse nurse cells turn over via a programmed cell death pathway closely resembling that used by Drosophila nurse cells. Somatic cells expressing V-ATPases and cathepsins associate with and acidify remnant nurse cells. Disrupting somatic acidification with a V-ATPase inhibitor in cultured ovaries blocks nurse cell programmed cell death (PCD). Our studies further illuminate a set of about 200 genes whose mRNAs accumulate selectively in dictyate oocytes as primordial follicles initially form. These include two tubulin gene transcripts, Tuba1c and Tubb2b, that promote Balbiani body development. Thus, mammalian oocytes are produced based on a conserved program shared by diverse animals that involves specification of nurse cells as well as oocytes within germline cysts.

## Results

### Identification of nurse cells based on reduced size and UMI content

Previously, we noticed that within ovarian germline cysts, germ cells in the process of cytoplasm and organelle transfer, putative nurse cells, are smaller in size than transfer recipients, putative oocytes (Figure 1A; Lei and Spradling, 2016). Oocytes use the transferred organelles, including centrosomes, to generate a Balbiani body (Figure 1B). Consequently, we re-analyzed data from a large scRNAseq analysis of mouse ovaries comprising seven time points spanning early cyst formation (E11.5) until the completion of cyst breakdown and primordial follicle formation (P5) (Niu and Spradling, 2020). Using Serualt 2, we re-clustered nearly 1,800 germ cells with a higher degree of segmentation, identifying 22 separate groups, rather than the previous 8 (Figure 1C; see Table S1-S2). Fifteen clusters comprising 75% of total cells had moderate to high levels of UMI/cell (Figure 1D, above dashed line) and localized on the outer circumference of the tSNE plot.

Using the sample ages and stage-specific meiotic gene expression data reported previously (Niu and Spradling, 2020), we observed that during normal perinatal development, female germ cell developmental states move in order around the tSNE periphery beginning with cluster 0a (pre-meiotic) at E11.5-E12.5 and ending with cluster 4b (P5 dictyate oocytes) (Figure 1C, Figure S1). In this higher resolution analysis, each meiotic stage is present mostly at one time point (Figure 1E,Table S2) and is separated into 2-3 ordered sub-clusters. The average UMI/germ cell varied with stage (Figure 1F) and increased significantly after birth in a similar manner to previous germ cell volume measurements, (Lei and Spradling, 2016; Figure 1G).

In contrast, seven clusters comprising 25% of cells (1N, 1Nr, 2Nr, 3N, 4N, 4Nr, 5) had moderately (1N, 3N, 4N) or strongly reduced (1Nr, 2Nr, 4Nr, 5) UMI/cell (Figure 1D, below dashed line) and distributed near the center of the plot. Six reduced UMI clusters were as stage-specific as the normal germ cells, and diverged from the main progression within specific meiotic stages (Figure 1E). These cells seemed likely to be nurse cells due to their high abundance and persistence, but they also could be cells or cysts undergoing apoptosis or fetal oocyte attrition. Cluster 5 cells (<3.5% of germ cells) displayed low UMI, high mitochondrial expression and derived from all time points; they were likely artifacts and not considered further. We carried out additional experiments to further document stage-specific low-UMI cells as nurse cells.

### Small germ cells act as nurse cells within cysts

If low UMI germ cells are nurse cells in the process of transferring their cytoplasmic contents to sister cells, then they should be located within germline cysts that also contain normally developing germ cells. To investigate this we carried out lineage labeling of E10.5 germ cells using very low levels of tamoxifen injection (Lei and Spradling, 2013). This procedure labels individual genetic clones consisting of several related cysts that are located close to each other in the ovary (generated by fragmentation of a single parental germline cyst). The results strongly supported the idea that the low UMI cells are cells that are transferring cytoplasm within cysts. For example, when an ovary lineage-labeled with YFP at E10.5 was analyzed at E16.5, a clone of 13 cells was labeled (Figure 2A, E16.5). By analyzing the DAPI channel it could be seen that one of the labeled cells was small, and consisted mostly of a nucleus, as expected if it was a member of cluster 2Nr. Following analysis at E18.5 a cyst was observed with two small cells with reduced cytoplasm (Figure 2A) suggestive in this case of membership in cluster 3N. Similarly, a lineage-labeled ovary analyzed at P1 revealed a two-cell cyst, where one daughter was small and only retained a rim of cytoplasm, suggest that it was a nurse cell that belonged to cluster 4Nr (Figure 2A, P1). In a similar manner, we analyzed clones between E14.5-P1 (Figure 2B), comprising more than 140 cysts (Figure 2C, red dots) and found that they contained 12.5%-36.8 small cells (Figure 2C, blue dots). The results showed that the nearly all the small cells are found within cysts; the turnover of small cells and generation of new ones from full sized cells that start to transfer cytoplasm causes the percentage of nurse cells to increase with time (see Figure 2D).

**Figure 2.**
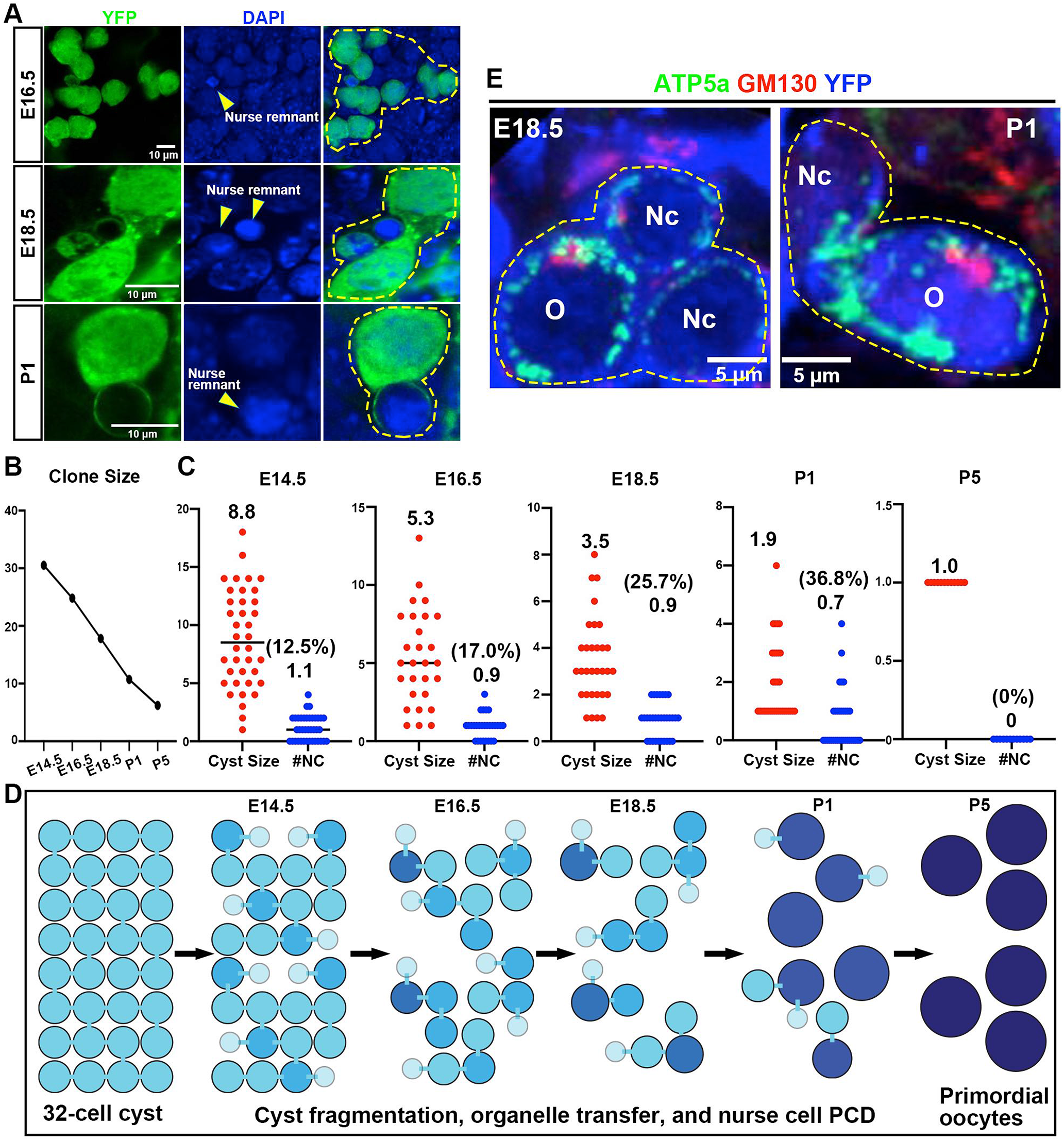
Germ cell cysts contain nurse cells with small size and low UMI. A. Individual germ cell clones (yellow dashed line) lineage-labeled with YFP (green) are shown from ovaries of the indicated ages. Germ cell(s) of reduced size (“nurse remnant,” yellow arrowheads) and size bars are indicated. B. Mean clone size decline vs developmental age due to cell turnover is plotted for clones induced in E10.5 germ cells. C. The number of to total cells and nurse cells is shown for cysts at E14.5, E16.5, E18.5, P1 and P5. At each age, the number of cells in many individual cysts is plotted (red dots), with the average number given above. The number of reduced-size germ cells per cyst is also (blue dots). The percentage of nurse cells per cyst is shown in parentheses. D. A model of showing how nurse cell transfer and PCD during E14.5 to P5 progressively reduces average cyst size and increases the fraction of nurse cells (small circles) per cyst. E. Examples of lineage-marked (YFP, blue) cysts at E18.5 and P1 illustrating organelle transfer between nurse cells (NC) and oocyte (O). ATP5a (mitochondria, green) GM130 (red, Golgi). Further analysis of these cysts is shown in Figure S2.

The high frequency of reduced-UMI germ cells detected by scRNAseq and observed within marked cysts is consistent with their function as nurse cells, but inconsistent with apoptosis of defective cells. The steady reduction in germ cell numbers during the nine days between E14.5 and P4 of about 80% (Figure 2B), would require 2.32 half-lives, suggesting an average lifetime of 3.9 days. Some of this time will be spent as full-sized intermediate cells in the process of trans-shipping cytoplasm between a low UMI cell and its final pro-oocyte destination. Low UMI cells themselves constitute about 24% of total cells (Table S2), suggesting that their average lifetime after beginning to lose cytoplasm is approximately 2.2 days (2.2/9 = 24%). This is consistent with our observation that the majority of cells in a given low UMI cluster are detected at a single time point, with much smaller proportions at the neighboring time points (Table S2).

The idea that the smaller cells are involved in transfer was further supported by labeling Golgi elements (GM130) and mitochondria (ATP5a) in lineage-labeled cysts (YFP) at E18.5 (Figure 2E, E18.5). A three cell cyst contains a larger cell (O) with most of the organelles, adjacent to two smaller presumptive nurse cells (NC) containing relatively few organelles but sharing the lineage marker. At P1 a YFP-labeled two-cell cyst shows a large organelle-rich oocyte, still attached to a nurse cell that has transferred nearly all its cytoplasm. The only remaining organelles are at the junction between the two cells (Figure 2E, P1). We conclude that the low UMI germ cells represent nurse cells in the process of transferring cytoplasm to other cyst germ cells. Thus, our data reveals nurse cell gene expression at four different stages of development (E14.5, E16.5, E18.5 and P1) as well as at earlier (1N, 3N, 4N) or more advanced states of transfer (1Nr, 2Nr, 4Nr).

### Nurse cell transcriptomes resemble those of their sister pro-oocytes

If nurse cells represent a distinct cell type, then their gene expression might significantly diverge from normal meiotic germ cells, as suggested by their separation into distinct clusters on the tSNE plot. However, this was not the case with 28 major meiotic-stage-enriched transcripts (Figure 3A). Expression of these genes within the reduced UMI germ cells (Figure 3A, right columns) largely follows the developmental pattern of normal germ cells of the same age (Figure 3A, left columns). For example, clusters 1N and 1Nr are present in E12.5-E14.5 ovaries, and are located on the tSNE plot (Figure 1C) near meiotic clusters 1a-1e (leptotene, zygotene). Figure 3A shows that clusters 1N and 1Nr express genes also expressed in normal leptotene and/or zygotene germ cells, such as Stra8, Tuba3a and ccnd1. They do not express premeiotic genes (Hmgn5, Hist1h2ap) or genes from later meiotic stages (such as the pachytene gene Sycp2, diplotene gene Uba52, or the dictyate genes Sohlh1). This is as expected if they arise from normal leptotene or zygotene cells that begin to transfer their cytoplasm to other germ cells within their cyst, changing their RNA profiles only slightly in the process. Similarly, the highly reduced UMI cluster 2Nr represents mostly E16.5 cells like the E16.5-specific pachytene clusters 2a-2c. 2Nr cells express all the genes in Figure 3A that are strongly expressed in pachytene, but show low or absent expression of the earlier or later expressed genes that are not expressed in pachytene (e.g. Rhox9, Phlda1, Sohlh1, Padi6). The E18.5 low UMI cluster 3N shows the same logic of expression. This cluster expresses diplotene genes such as Uba52, Syce3 and Id1, but not genes whose expression is absent in diplotene such as Stra8 or Rec8. Finally, the P1-labeled low UMI 4N and 4Nr also preferentially express normal dictyate genes from among the genes in Figure 3A. We confirmed these conclusions by comparing up-regulated and down-regulated genes generally during successive timepoints for normal meiotic genes (Figure 3B-D) to the changes in the corresponding nurse cells (Figures 3B’-3D’). The major genes changing in either direction are the same in normal meiotic cells and in nurse cells.

**Figure 3.**
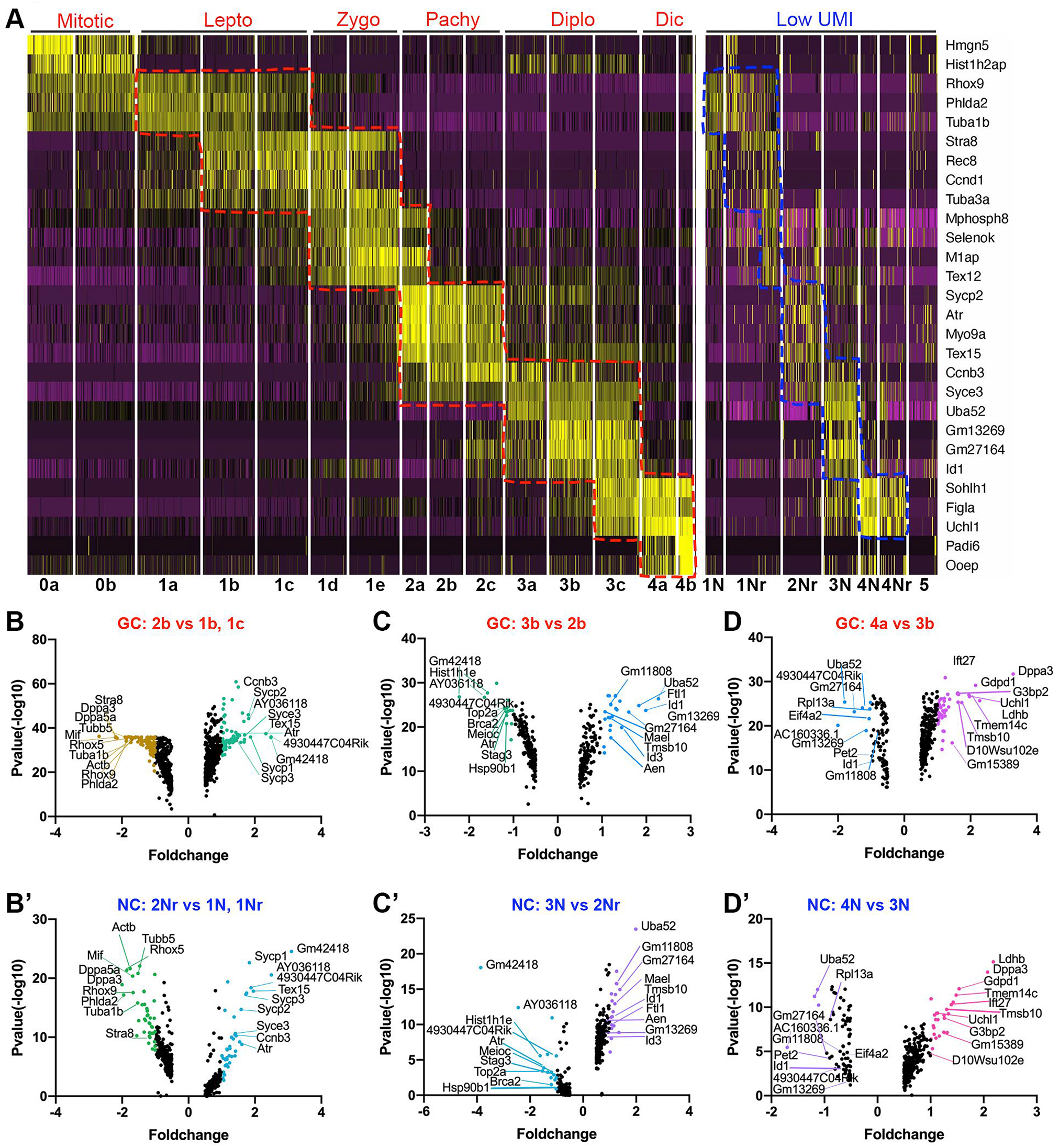
Full size meiotic germ cells and nurse cells express similar transcripts. A. The relative expression of the genes indicated at right within the indicated scRNAseq clusters (bottom) which are listed in their order of meiotic stage (cluster number). Yellow indicates high and black low expression. Peak expression (yellow) in full-sized germ cells (left columns) and in lowUMI nurse cells (right columns) is very similar at the corresponding stages. B.-D. Volcano plots showing gene expression changes between successive stages for full sized meiotic germ cells (GC, red) (B,C,D) and nurse cells (NC, blue) (B’, C’ and D’). The strong similarities in gene expression changes in GC and NC are indicated by the identification of major downregulated (left) and upregulated (right) genes in each plot, which correspond closely.

We searched for changes in less abundant genes that might be responsible for the divergence of normal meiotic cells and nurse cells on tSNE plots. In particular, we were interested in common changes that occur in multiple nurse cell groups at different times during meiosis. We identified a number of genes that are upregulated in nurse cells compared to the meiotic germ cells from which they derived. Some of these may be due to delayed mRNA turnover in nurse cells. For example, genes like Stra8 that are highly expressed at leptotene-zygotene (E14.5) but sharply downregulated at pachytene (E16.5) show elevated expression in E16.5 nurse cells (2Nr). However, we also identified about 300 genes upregulated in nurse cells at E14.5 and E16.5 (see Table S4). GO analysis of these genes included: “response to estradiol (GO:0032355) FDR: 8.47E-3,” “response to BMP (GO:0071772) FDR: 8.20E-3”, and “positive regulation of programmed cell death (GO:0043068) FDR: 3.27E-3,” suggesting that nurse do respond to intercellular signals and alter their gene expression.

### Somatic cells and the cyst cytoskeleton contribute to oocyte development

The strong similarity in gene expression between meiotic germ cells and nurse cells, suggested that nurse cells develop as normal meiotic cells, but at varying times during meiosis they receive an external signal to start transferring their cytoplasm and organelles within the cyst. One possible source for such as signal is the somatic pre-granulosa cells that surround cysts of developing germ cells throughout their transition from forming cyst to completed follicle (Figure 4A). To investigate the role of the epithelial pre-granulosa (EPG) cells that surround developing wave 2 follicles that will form the primary reserve, we ablated EPGs as described previously using Lgr5-DTR-EGFP mice and diptheria toxin injection at E14.5 (Figure 4B; see Niu and Spradling, 2020). Ablation was highly effective, as the number of Lgr5+ cells, which normally surround developing germ cells (Figure 4B) was greatly reduced. Germ cells continued to develop although their numbers become significantly lower after E18.5. Golgi elements decreased in P1 (Figure 4C) and P4 (Figure 4D) germ cells and did not cluster into Balbiani body. Not only Golgi/per cell (Figure 4E), but mitochondria/per cell (Figure 4F) and Pericentrin (centrosomes)/per cell (Figure 4G) decreased. Thus, germ cells lacking interactions with EPGs did not undergo the normal, large increase in organelle numbers after E18.5, and did not form Balbiani bodies.

**Figure 4.**
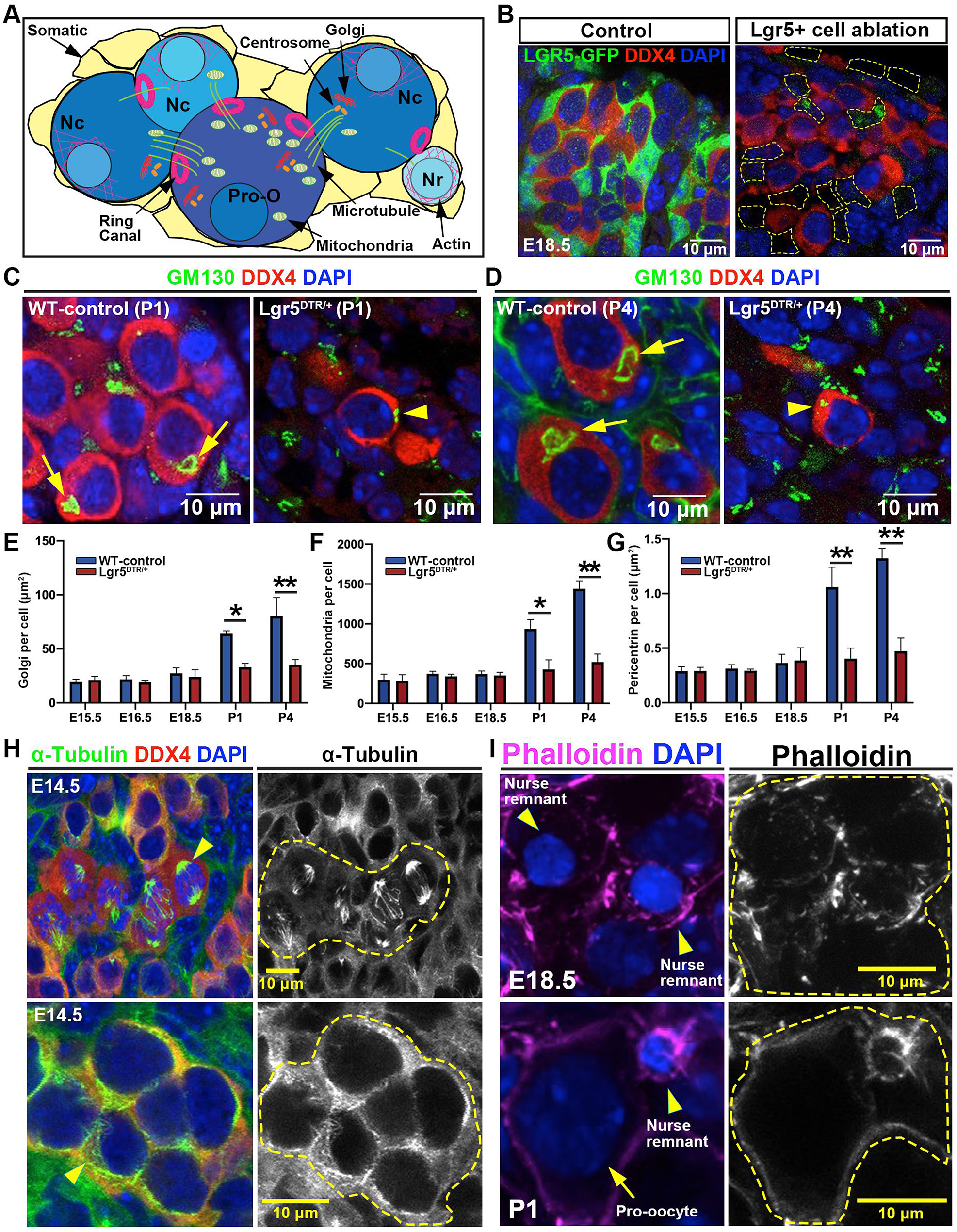
Somatic cell signals and the cyst cytoskeleton guide nurse cell and oocyte selection. A. A model depicting a five-cell cyst containing a central oocyte with 3 intercellular bridges (IBs) that will become the oocyte, and 3 smaller-sized nurse cells (Nc) and 1 nurse remnant (Nr), each with 2 or 1 IBs, and surrounded by somatic cells (tan). The location of IBs (pink circles), microtubules (green lines), actin fibers (purple lines), Golgi elements (long red ovals), mitochondria (green ovals), and centrosomes (orange small circles) are shown. B-G. Effects of granulosa cell ablation on germ cell organelles. B. Lgr5-expressing pre-granulosa cells (green) surround germ cells expressing DDX4 (red) at E18.5. Following Lgr5-Dtr-EGFP mediated EPG cell ablation (see Methods), Lgr5-expressing EPGs (control) are gone (ablation, yellow dashed boxes). C-D. Control and ablated ovarian cells showing that ablation greatly reduces the size and structure of Bb-associated Golgi (green, arrows) at P1 (C) and P4 (D). E-G. Bar graphs showing the Golgi/cell (E), mitochondria/cell (F) and Pericentrin (centrosomes)/cell (G) in ovarian cells from control (blue) and ablated (red) animals. H-I. The cyst cytoskeleton. H. alpha-Tubulin immunofluorescence of E14.5 ovarian cells showing the last synchronous cyst metaphase (upper panels) and the rich microtubule cytoskeleton seen throughout cells in the cyst (lower panels). The right panels show the tubulin channel alone. I. E18.5 (upper) and P1 (lower) ovarian cysts stained with phalloidin to reveal actin fibers in germ cells. Large aggregations of actin are seen associated with small nurse cell nuclei (nurse remnants, yellow arrowheads) whereas no similar actin staining is associated with full-sized germ cell or oocyte nuclei. Size bars are indicated.

Another well-established contributor to oocyte development during the cyst stage in invertebrates is the cyst cytoskeleton (Figure 4A). We labeled cyst microtubules with an anti-alpha-tubulin antibody, and observed synchronous cyst mitoses up to E14.5 (Figure 4H, upper panels). Other cysts of the same age that had completed mitotic divisions retained abundant cytoplasmic microtubules (Figure 4H, lower panels). While the degree of microtubule asymmetry in the cyst remains to be determined, alterations to this cytoskeleton may explain why treatment of cultured ovaries with low levels of anti-microtubule drugs impairs the formation of primordial oocytes with normal Balbiani bodies (Koch and Spitzer, 1983; Lei and Spradling, 2016).

The actin cytoskeleton plays critical roles during Drosophila cyst development. Bundled actin fibers form in stage 10A that attach to nurse cell nuclei just before bulk cytoplasmic transfer at stage 10B. If these fibers are disrupted by mutation in profilin, or a variety of actin bundling proteins, nuclei move toward the oocyte and block transfer (reviewed in Hudson and Cooley, 2002). We looked for the presence of actin fiber in mouse cysts before and during transfer. Strong focal accumulations of actin were observed surrounding nurse cell remnant nuclei in cells that were transferring cytoplasm but not nuclei (Figure 4I).

### Programmed cell death in mouse nurse cells involves cell acidification

Another nurse cell process that may have been conserved during evolution was addressed by examining mouse nurse cell programmed cell death (PCD) (see Figure 5A). This process has been analyzed in depth during nurse cell “dumping” and PCD in the Drosophila ovary (reviewed in Lebo and McCall, 2021). A major indicator of the somatic-driven PCD pathway in the Drosophila ovary is nurse cell acidification (Mondragon et al. 2019), which is driven by fusion with acidic vesicles generated within surrounding somatic cells. We used the acidophilic dye lysotracker and observed many germ cells with smaller than average nuclei become Lysotracker positive from E16.5-P1 (Figure 5B). Counts showed that between 10% and 33% of germ cells were lysotracker positive, with the peak at E18.5 (Figure 5C). The small nurse cells undergoing acidification are surrounded by Lgr5-positive cells, a marker for pregranulosa cells (Figure 5D).

**Figure 5.**
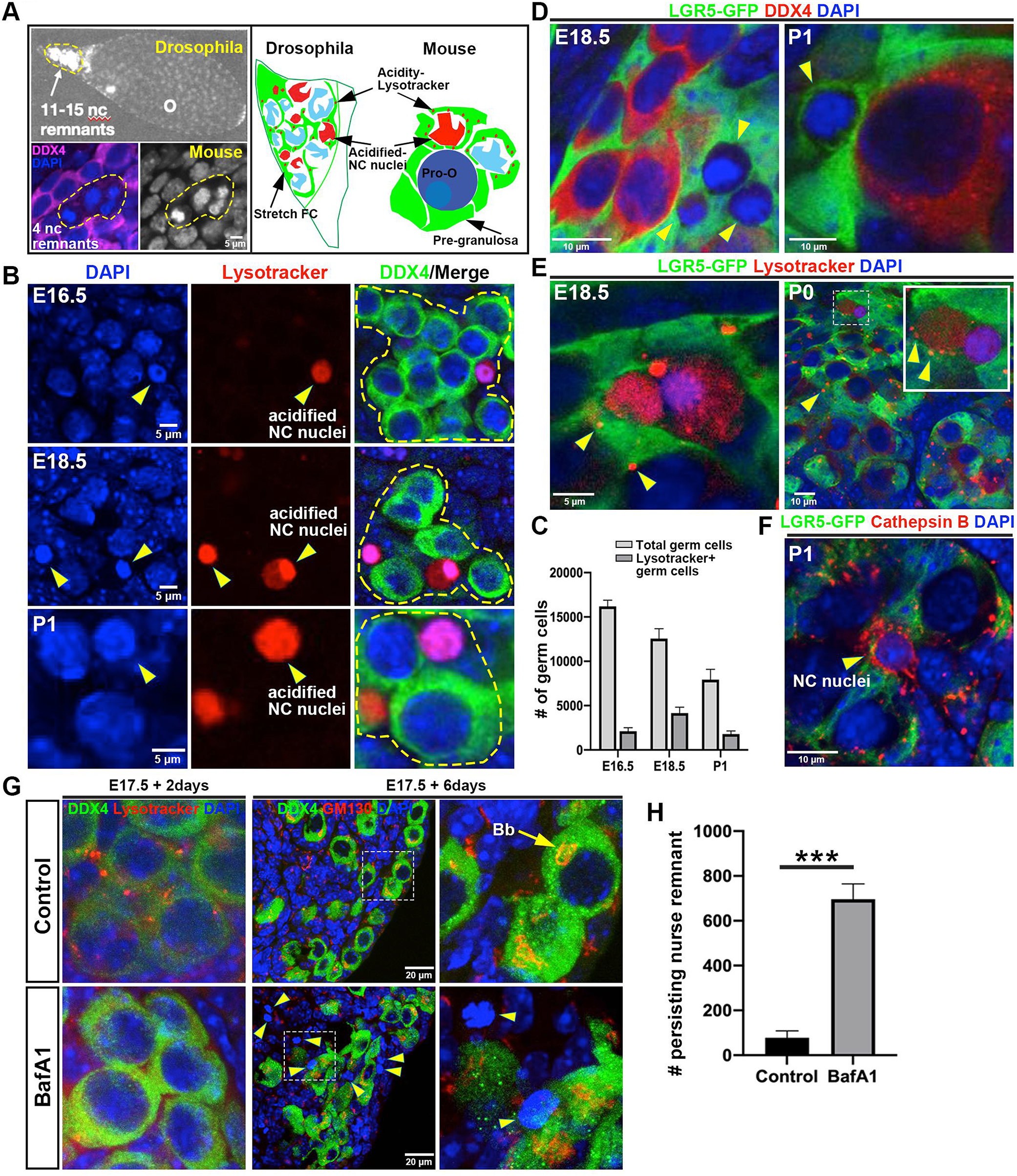
Programmed cell death of mouse nurse cells. A. Left: 11-15 Drosophila (upper) and 4 mouse (lower) nurse remnants showing their persistence and association. Right: Drawings showing the acidification (Lysotracker, red) of Drosophila NC surrounded by stretch follicle cells, and of mouse NC remnants surrounded by pre-granulosa cells. B. Acidification of mouse nurse cells is revealed by positive Lysotracker staining (red) in a minority of small cyst cells at three time points. C. The number of total and Lysotracker-positive germ cells is shown; the percentage increases from about 10% at E16.5 to peak at about 33% at E18.5. D. Cyst nurse cells (yellow arrowheads) are surrounded by Lgr5+ pre-granulosa cells at E18.5 and P1. E. A subset of nurse cells positive for DDX4 (green, germ cell marker), are also positive for the acidification marker Lysotracker (red), at E16.5, and P0. F. Cathepsin B is expressed in some Lgr5+ EPG cells surrounding nurse cell nuclei at P1, but is not found at P4, when PCD is complete (not shown). G-H. Ovary culture in the absence (Control) or presence (BafA1) of the V-ATPase inhibitor Bafilomycin A beginning at E17.5. After 2 days, Lysotracker-positive vesicles are visible in control but not BafA1 treated cells. After 6 days, nurse cells have regressed and oocytes with Bbs have formed in control cultures, but in BafA1 cultures, many nurse cell remnants remain (yellow arrowheads). H. Quantitation of excess persisting nurse remnants remain after 6 days when V-ATPases are inhibited with BafA1. Size bars are indicated.

In Drosophila, somatic cell V-ATPases generate a pH gradient across the somatic cell membrane in the nurse cell and exocytosis machinery then transfers cathepsins (acid activatable proteases) to drive acidification and proteolysis non-autonomously. The Lgr5+ pre-granulosa cells surrounding mouse nurse cell express lysotracker positive vesicles during stages when nurse cell turnover is high (Figure 5E). Multiple mouse V-ATPases are active in these pre-granulosa cells as well a multiple cathepsins based on previous scRNAseq analyses (Figure S5; Niu and Spradling, 2020). The most highly expressed cathepsin in granulosa cells, Ctsl, is the ortholog of the Drosophila-capthesin Cp1 which is required for nurse cell turnover (Mondragon et al. 2019), while Cathespin B (Ctsb) is the 2nd most highly expressed and is found in pregranulosa cells surrounding remnant nurse cells (Figure 5F). This expression, like nurse cell turnover, is gone by P4 (Figure S5).

To investigate if acidification is required for mouse nurse cell turn over, we cultured ovaries in vitro starting at E17.5 either in the presence or absence of the vacuolar ATPase inhibitor Bafilomycin (BafA1). After 2 days without the inhibitor, oocyte maturation into primordial follicles is underway (Figure 5G) and is largely complete after 6 days in culture as indicated by the presence of GM130-positive Balbiani bodies (Figure 5G, arrows). In contrast, in the BafA1-treated cultures, lysotracker vesicles are no longer seen in somatic cells, and many persistent nurse cell nuclear remnants remain (Figure 5G, arrowheads). Quantitating this effect showed that disruption of V-ATPase function with BafA1 strongly inhibited nurse cell PCD and generated a massive increase in nurse cell nuclear remnants (Figure 5H).

### Pro-oocyte enriched RNAs include two tubulin genes requred for Balbiani body formation

Female germ cell gene expression during E11.5-P5 has been extensively studied using both developmental genetic (Wang et al. 2020) and high throughput studies (Soh et al., 2015; Miyauchi et al. 2017; Kojima et al., 2019; Nagaoka et al. (2020), Niu and Spradling, 2020). Figla (Joshi et al. 2008; Wang et al. 2020), and Taf4B (Grieve et al. 2014; Grieve et al. 2016) are required for normal meiotic gene expression, and Figla also promotes oocyte growth and development, acting in part by regulating transcription factors such as Nobox, Lhx8, Sohlh1 and Sohlh2 (Liang et al. 2007; Joshi et al. 2008; Wang et al 2020). We identified nearly 200 genes that are upregulated in dictyate oocytes, and include many target genes in this cascade (Figure S6, Table S5). Only 13% of these genes were expressed in nurse cells at even 50% of their level in oocytes, while a majority were expressed at less than 10% of oocyte levels. Likewise, genes transcripts encoding members of the subcortical maternal complex (Li et al. 2010) were present in P1 nurse cells at 0.00-0.46 the level in oocytes, while immunostaining of NLRP5 protein was detected preferentially in oocytes (Figure S6F). In addition, we identified a set of 175 genes that were enriched in normal-sized germ cells at multiple stages but under-expressed in nurse cells (Figure S2).

Two tubulin genes, Tuba1c and Tubb2b fall into the oocyte-enriched class (Figure 6A). Germline loss of Tuba1c prevented the final assembly of the Golgi elements in the P1 oocyte into a mature Balbiani body by P4 (Figure 6B) in about 40% of oocytes (Figure 6F). This same defect become even more frequent (55%) in the double mutant of both these genes (Figure 6C, F). A model of the dispersed centrosome in P1 oocytes (Figure 6D, left) and how centrosome aggregation converts separate organelles into a completed Balbiani body by forming a new microtubule organizing center is shown (Figure 6D, right). These changes are frequently disrupted in the absence of Tuba1c and Tubb2b (Figure 6E). Further analysis (Figure S6) showed that in addition to Golgi elements, cell volume, mitochondria/cell and pericentrin per cell were reduced by about 25%, suggesting that organelle transfer prior to oocyte completion is also affected..

**Figure 6.**
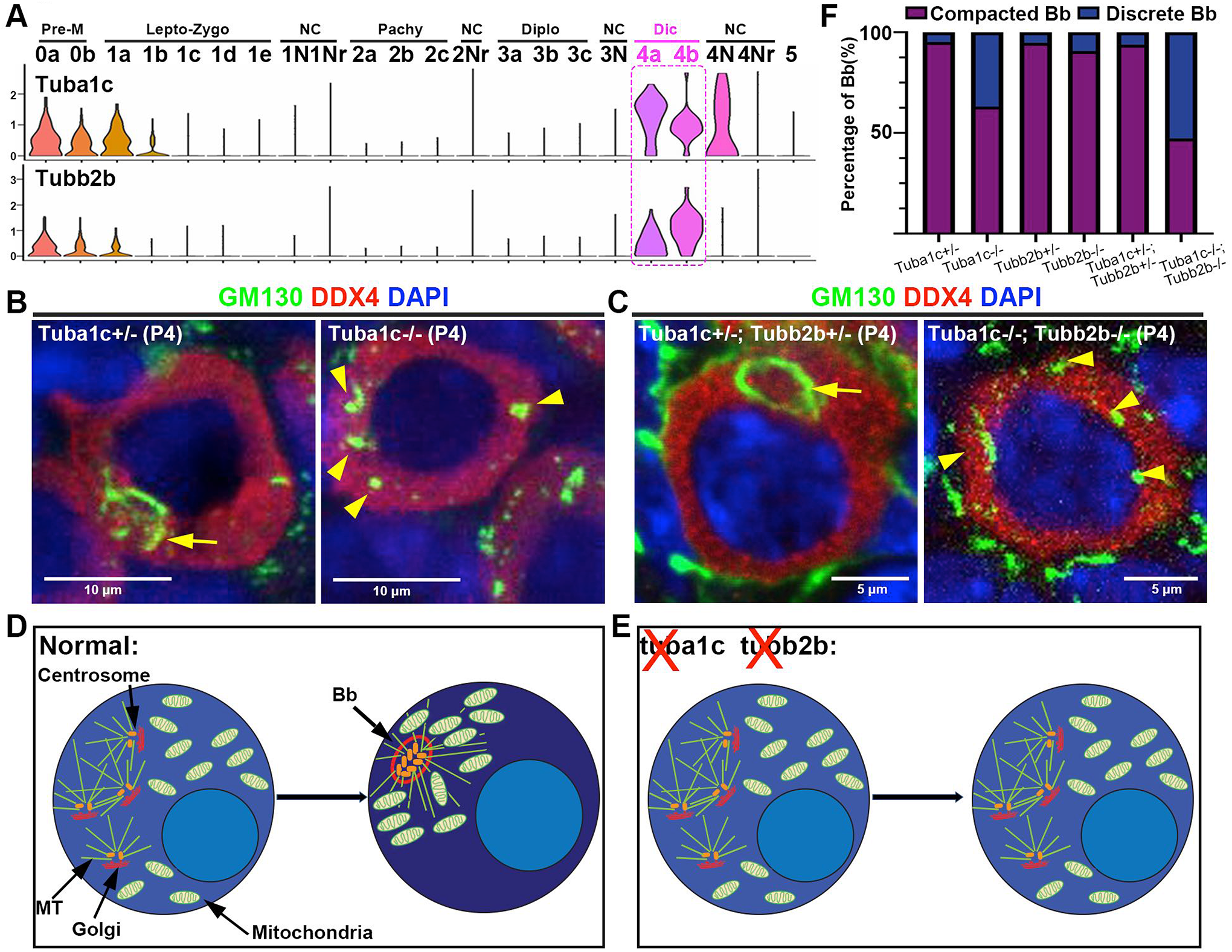
Tuba1c and Tubb2b contribute to Balbiani body formation in dictyate oocytes. A-C. scRNAseq plot shows that Tuba1c and Tubb2b are induced in dictyate oocytes (A). B,C. P4 oocytes are shown with either a normal Balbiani body in control Tuba1c −/+ heterozygotes, or with an abnormal Bb, where discrete Golgi elements (GM130, green) failed to reorganize into a compact structure by P4, a defect seen in about 40% of Tuba1c (−/−) oocytes (B,F). A higher frequency of abnormal Bb formation, 55%, is seen in Tuba1c (−/−); Tubb2b (−/−) oocytes (C, F). D-E. Model for role of tubulin in Balbiani body morphogenesis. D. Left cell: 4 Golgi elements in concert with centrosomes have arrived in a P1 oocyte following transfer from nurse cells (not shown). By P4 (right cell), with the assistance of Tuba1c and Tubb2b, the centrosomes aggregate and nucleate new microtubule arrays (green lines) which allow Golgi elements to form a shell around the centrosomal core. E. In genotypes lacking Tuba1c and enhanced by the absence of Tubb2b, nurse cell transfer still occurs (left cell), but centrosomes fail to aggregate and generate a new MT array to complete re-assembly of organelles into a compact Balbiani body.

## Discussion

### Mouse oocytes develop with the assistance of nurse cells in germline cysts

Previous studies of mouse fetal germ cell development within germline cysts identified smaller cyst cells that transfer cytoplasm and organelles into larger presumptive oocytes (Pepling and Spradling, 2001; Lei and Spradling, 2016). Here, we were able to identify and quantitate these presumptive nurse cells based on their small size and reduced UMI/cell. More than 25% of total germ cells in E14.5 to P5 mouse ovaries are active nurse cells that reside within germline cysts as sisters to the ~20% of cells that will become oocytes. Before being selected, germ cells that will become activated as nurse cells enter meiosis and pass through prophase stages like their sister germ cells. Regardless of their cell cycle stage, once selected nurse cells transfer their cytoplasm, organelles but not their nuclei into either an intermediate nurse cell, or directly into a pro-oocyte over a period of about 2.2 days. During transfer they gradually become depleted cell remnants, containing little more than a nucleus and a tiny rim of cytoplasm, but still surrounded by pre-granulosa cells, which go on to mediate their final turnover. Thus, under favorable conditions, the large number of female germ cells that die during perinatal stages are not due the culling of defective oocytes with chromosome abnormalities (Tilly, 2001), mitochondrial dysfunction (Palozzi et al. 2018), or LINE element activation (Tharp et al. 2020), but due to nurse cells turning over according to a developmental program that contributes to oocyte production. However, it remains possible that these other processes account for a small fraction of oocyte turnover.

### Three mechanisms likely help specify particular cyst cells to be oocytes or nurse cells

How oocytes are selected among cyst cells has remained only partially understood despite decades of research in other animals. The relatively simple, invariant cysts found in Drosophila contain 16-cells interconnected in a maximally branched pattern, containing a polarized microtubule-based cytoskeleton whose minus ends all point to the initial cell. This cell eventually becomes the only oocyte, after organelles and unknown “determinants” are transported into it, while the remaining 15 cells develop as nurse cells (King, 1970; deCuevas and Spradling, 1998; Matova and Cooley, 2001; Roper and Brown, 2004; Huynh and St.Johnston, 2004). Mouse oocytes are chosen within even more complex 32-cell initial cysts, that specify about 6 oocytes and 26 nurse cells. Our studies suggest that many of the same mechanisms involved in Drosophila oocyte/nurse cell specification also act on mouse cyst cells. We observed that a robust microtubule cytoskeleton that could influence the movement of cytoplasm undergoing transfer has developed in mouse cyts by shortly after their final cyst mitotic division at E14.5 (Figure 4H). Ablation of somatic EPG cells beginning at this time, blocked the transfer of Golgi, mitochondria and centrosomes into downstream cells (Figure 4 B-G). Finally, studies reported elsewhere argue that an intercellular bridge-dependent signal locally represses cytoplamic transfer (Ikami et al. 2021). In mouse, the combined action of these mechanisms must control cyst fragmentation, nurse cell/oocyte specification within multiple cyst fragments, and the time individual nurse cells begin to transfer cytoplasm throughout the 9 day interval between E14.5 and P5.

### Mouse nurse cells resemble pro-oocytes until signaled to transfer their cytoplasm and organelles

The ability to identify nurse cells before they have transferred most of their cytoplasm to an oocyte, allowed us to test whether nurse cells begin to differentiate in the early cyst into a distinct cell type. All the nurse cells within each mouse cyst do not transfer cytoplasm at the same time, but instead seem to become active from the edges of the cyst toward the center (Lei and Spradling, 2013). We compared nurse cells activated for transfer at E14.5, shortly after cyst completion, or only at P1, when few total nurse cells remain. We found that all full sized cells continued to express very similar transcriptomes up until the time a nurse cell begins to undergo transfer and decrease in size (Figure 3). These findings provide a molecular validation of the histological similarity observed previously in studies of meiotic fetal germ cells. However, more detailed analysis of genes enriched in nurse cells at E14.5 and E16.5 showed that important differences in gene expression do eventually arise in nurse cells, including the upregulation of genes that promote programmed cell death. Moreover, nurse cell enriched gene expression suggested that BMP and estradiol signals, presumably from ovarian somatic cells, participate in activating nurse cell cytoplasmic transfer.

### High conservation of an ancient nurse cell program

We observed striking similarities in the formation of germline cysts between mouse and Drosophila and in the potential mechanisms that likely select oocytes and nurse cells. The manner in which nurse cell remnants eventually turnover was also highly conserved. The long duration of mouse nurse cell remnants, their accumulation in small groups, and their turnover by a somatic cell-controlled process of programmed cell death involving acidification and cathepsin action, were all very similar to the well characterized Drosophila nurse cell PCD program (Lebo and McCall, 2021). The fact that nurse cell turnover is likely to be highly conserved, and does not follow the canonical apoptosis pathway, raises questions about some previous studies of cyst cell fates. The conclusions that all Xenopus cyst cells become oocytes (Kloc et al. 2004), and that most cyst cells in fish likely develop as oocytes (Jamieson-Lucy and Mullins, 2019) are largely based on a failure to find evidence for apoptosis as cyst cells break apart. It would be worthwhile to look for evidence of acidification of a fraction of cyst germ cells in these systems. Ultimately, lineage tracing will be needed to prove what fraction of cyst cells become oocytes.

Once the nurse cell program is completed as primordial follicles prepare to form, the conservation between mouse and Drosophila appears to end. The dramatic induction of about 200 genes in dictyate mouse oocytes does not have a currently recognized counterpart in Drosophila oogenesis at the time of follicle formation. This may represent a point where mouse oocytes differentiate but where in the evolutionarily revised scheme of oogenesis developed by higher insects, nurse cells now differentiate instead (Deluca and Spradling, 2020).

### What is the advantage to oocytes of developing within cysts assisted by nurse cells?

Ideas about the importance of nurse cells have been strongly influenced by observations on the massive nurse cells that are present in Drosophila and many other holometabolus insects. Since these cells are responsible for synthesizing the great bulk of the oocyte cytoplasm, they have been widely viewed as helping the oocyte grow to a large size which in most animals represents the largest body cell. Producing the massive oocytes found in Drosophila and many insects would be a very slow process if growth was supported only by the oocyte’s 4c nucleus.

However, it is much less clear that the early growth of the mouse or Drosophila oocyte prior to follicle formation requires the assistance of nurse cells. Mouse oocytes do not significantly increase in volume until after E18.5. While oocytes do grow 4-fold between E18.5 and P5, this increase may not be caused by nurse cell cytoplasmic transfer. The new transcriptional program induced in diplotene oocytes by Figla and its downstream targets (Wang et al. 2020), might be sufficient stimulate the observed oocyte growth. It might be easier to generate oocytes of a uniform size using a growth program regulated by feedback, compared to relying on cytoplasmic transfer from a potentially variable number of nurse cells.

There are several reasons other than growth stimulation that female gametes might benefit from sharing cytoplasm with their sister germ cells. The first would be to obtain the same advantage cysts are thought to provide for male gamete development- protection against selfish meiotic drive elements and other parasites. While female cyst cells remain diploid, unlike in males, cytoplasm sharing might interfere with selfish elements whose gene products act on chromosomes to bias their positioning on the meiotic I spindle. Cytoplasmic sharing may also help neutralize newly acquired or newly activated transposons that stimulate strong piRNA production in only a few cyst germ cells. Sharing such piRNAs has the potential to jumpstart the repression of these elements in all cyst cells.

The final potential advantage of cysts and cytoplasmic transfer is their role in generating the Balbiani body. Although the term “Balbiani body” has been applied to a diverse set of oocyte structures, we propose that the transfer of organelles from nurse cells into future oocytes at the time of follicle formation gives rise to a fundamental type of Bb that contributes to oocyte patterning. In particular, the multiple centrosomes acquired by transfer aggregate into a single cytoplasmic focus in dictyate oocytes that may reorient microtubules and drive Balbiani body morphogenesis by assembling Golgi elements into a shell and concentrating mitochondria peripherally (Figure 6D). Generating a cellular signaling center by collecting centrosomes from interconnected sister cells may be an ancient mechanism of oocyte polarization. In Drosophila, centrosomes transferred from nurse cells leave the Balbiani body as the follicle is forming, and migrate to the oocyte posterior along with oskar and CPEB mRNAs which may eventually establish this region as a signaling center (Mahowald and Strassheim, 1970; Cox and Spradling, 2003). Here we found that two tubulin genes, Tuba1c and Tubb2b, that are induced in dictyate oocyte, are important for mouse Bb assembly (Figure 6). At least one of these genes, Tubb2b, is developmentally regulated and functionally important in neural development (Bittermann et al. 2019).

### The conservative and radical evolution of female gametogenesis revealed by comparing mouse and Drosophila

Mouse female gametogenesis now joins Drosophila as one of the best understood examples of oocyte production; these systems share many features but also diverge sharply. Early germ cell development within cysts is found in essentially all animal groups in male germ lines, and has been retained in female germ lines within diverse animals including mouse and Drosophila, but not in all groups. Thus, evolutionary conservation likely explains the many conserved features of early oogenesis we highlighted here, including cyst formation with intercellular bridges, cyst cytoskeletal asymmetry, somatic regulation of oocyte development and nurse cell turnover, and nurse cell organelle transfer leading to the Balbiani body. This initial period of female gamete production includes key developmental features such as parasite protection, sex determination, meiosis and recombination, oocyte polarization and follicle formation that may be responsible for the high conservation.

However, our studies also brought to light at least two major evolutionary changes to this ancient system of female gametogenesis. Drosophila and most holometabolous insects beginning about 250 Myr ago greatly accelerated the speed of oogenesis by suppressing nurse cell breakdown, amplifying nurse cell genomes by polyploidization, and employing nurse cells to synthesize almost all the oocyte contents. Mouse and eutherian mammals, in contrast, about 120 Myr ago limited oocyte growth, diminished vitellogenesis and evolved an elaborate placenta and implantation system allowing the vastly greater resources of the mother to nourish her developing embryos. Both events impacted the conserved events spanning cyst to follicle formation. Keeping nurse cells required their turnover to be postponed, and the packaging in follicles of all cyst cells. Another potential divergence between mouse and Drosophila is the dramatic upregulation of genes controlled by newly induced transcription factors in dictyate mouse oocytes (reviewed in Wu and Dean 2020). No corresponding induction of oocyte genes has been found in early Drosophila oocytes, which soon enter meiotic arrest and form a karyosome. Instead, Drosophila nurse cells differentiate as somatic cells (Deluca and Spradling, 2020). They subsequently join other ovarian somatic cells in regulating oocyte growth, nurse cell breakdown, and ovulation using potentially conserved hormonal (insulin, steroid hormones) and inter-tissue coordination. Thus, these studies are clarifying not only how mammalian oocytes arise, but also the evolution of female gametogenesis, due to the recognition by germ cell researchers that studying invertebrate as well as vertebrate animals greatly accelerates the advancement of biomedical knowledge.

## Methods

### Animals

Mouse experiments in this study were performed in accordance with protocols approved by the Institutional Animal Care and Use Committee (IACUC) of the Carnegie Institution of Washington. CAG-creER (004682), Lgr5-CreERT2 mice (008875), and R26R-EYFP reporter mice (006148) were acquired from the Jackson Laboratory. Tuba1c^em1(IMPC)J^/Mmjax mice (051205-JAX), and Tubb2b^em1(IMPC)J^/Mmjax mice (046114-JAX), were acquired from Mutant Mouse Resource & Research Centers (MMRRC). Lgr5-DTR-EGFP mice were obtained from Genentech (South San Francisco, CA).

### Labeling and Tracing Experiments

The R26R-EYFP females were crossed with the CAG-creER males, those with a vaginal plug were considered as E0.5. The pregnant females at E10.5 were given a single intraperitoneal injection of tamoxifen (Tmx; 10 mg/ml in corn oil (Sigma)) at 0.2 mg per 40 g body weight

### Diphtheria Toxin injection

Pregnant mice (E14.5) were injected i.p. with 10 μg/kg of diphtheria-toxin solution (D0564, Sigma) in PBS.

### Immunofluorescence

Ovaries were fixed in cold 4% Paraformaldehyde overnight, incubated sequentially in 10% and 20% sucrose in PBS overnight, embedded in OCT, and stored at −80°C until cryosectioning. After high-temperature antigen retrieval with 0.01% sodium citrate buffer (pH 6.0), the frozen sections (10 μm) were blocked with 10% normal donkey serum for 30 mins, and then incubated with primary antibodies overnight at 4°C. The primary antibodies used are presented in Table S1. The sections were washed with wash buffer and incubated with the appropriate Alexa-Fluor-conjugated secondary antibodies (1:200, Invitrogen) at room temperature for 2 hs. After staining with DAPI, samples were analyzed using confocal microscopy (Leica SP5).

### Whole-mount staining of mouse fetal ovaries

Ovaries were fixed in cold 4% PFA for 4 hours. After washing in PBST_2_ (PBS with 0.1% Tween-20 and 0.5% Triton X-100) for 90 minutes, ovaries were then incubated with primary antibodies for 48 hours at 4°C. After washing in PBST_2_ for 90 minutes, ovaries were incubated with secondary antibodies overnight at 4°C. After washing in PBST_2_ (90 minutes) and DAPI staining (1 hour), samples were analyzed using confocal microscopy (Leica SP5).

### Ovary *in-vitro* culture

Ovaries were dissected in cold PBS, and then cultured in 1 mL DMEM/F12 medium (ThermoFisher) at 37°C in an atmosphere of 5% CO2. The medium was supplemented with antibiotics penicillin and streptomycin (Fisher) to prevent bacterial contamination. Cultured ovaries were carefully placed on a membrane insert (0.4 μm, Millipore) and treated with Bafilomycin A1 (BafA1, 100nM, Abcam), which is a selective (V)-ATPase inhibitor. After 2 or 6 days in culture, ovaries were fixed in 4% PFA for further analysis.

### LysoTracker staining of mouse fetal ovaries

Lysotracker red (L7528, Thermofisher) was diluted 1:10,000 in DMEM/F12 medium. Ovaries were carefully dissected and cultured *in-vitro* in 1 mL Lysotracker red added medium at 37°C for 12 hours. Ovaries were then harvested and fixed in 4% PFA for further analysis.

### Quantifying germ cell number, follicle number, nurse cell number

For germ cell or follicle counts, collected ovaries were fixed in 4% PFA, incubated in sucrose, embedded in OCT, and sectioned to a thickness of 10 μm. The sections were stained with DDX4 antibody and every fifth section was analyzed for the presence of germ cells. The cumulative germ cell or follicle counts were multiplied by five. For nurse cell counts, single germ cell was labeled at E10.5 through low TMX injection (see labeling and tracing experiments for details), and ovaries were harvested at different timepoints for analysis. Collected ovaries were fixed in 4% PFA and stained with DDX4 and EGFP antibodies for whole-mount staining. The number of nurse cells that are smaller in cell volume were counted in each cyst.

### Tissue dissociation and single cell library preparation

Ovaries were dissected and placed in 1X PBS on ice, then dissociated into single cells using 0.25% Trypsin at 37°C with pipet trituration at intervals. E12.5 ovaries were dissociated for 20 min, E14.5 ovaries for 40 min, E18.5 ovaries for 1h, P1 and P5 ovaries for 80 min. After adding 10% FBS, the dissociated cells were passed through 70 μm and 30 μm cell strainers, separately. About 10,000 live cells were then loaded per sample onto the 10x Genomics Chromium Single Cell system using the v2 or v3 chemistry per manufacturer’s instructions (Zheng et al., 2017). Single cell RNA capture and library preparations were performed according to manufacturer’s instructions. Sample libraries were sequenced on the NextSeq 500 (Illumina). Sequencing output was processed through the Cell Ranger 2.1.0 mkfastq and count pipelines using default parameters. Reads were quantified using the mouse reference index provided by 10x Genomics (refdata-cellranger-mm10v1.2.0). 10X genomics chemistry version 3’c2; mouse transcriptome mm10, cell ranger 2.1.1.

### Cell identification and clustering analysis

Filtered count matrices for each library were normalized as described in the package “Seurat” (Satija et al. 2015; https://satijalab.org). Batch correction was performed using the *JackStraw* and *RunCCA* functions in the Seurat package.

## Data Availability

Sequence data have been deposited in the GEO database (GSE136441).

## Acknowledgements

We thank the Carnegie mouse facility and its manager Dr. Eugenia Dikovsky, for outstanding support. We thank the Johns Hopkins University School of Medicine Biotechnology center for assistance with some of the scRNAseq experiments. We especially thank Allison Pinder and Fred Tan of the Carnegie Embryology Biotechnology Center for assistance in carrying out scRNAseq and in data analysis. We thank Dr. Frederic J. de Sauvage (Genentech, Inc.) for kindly providing us with Lgr5-DTR-EGFP mice. We thank Dr. Jurrien Dean (National Institute of Diabetes and Digestive and Kidney Diseases, NIH) for kindly providing us with NLRP5 antibody.

## Supplemental Online Materials

### Supplemental Tables

Table S1 Gene expression data for 22 clusters (mUMI/cell).

See Excel File S1.

**Table S2.**
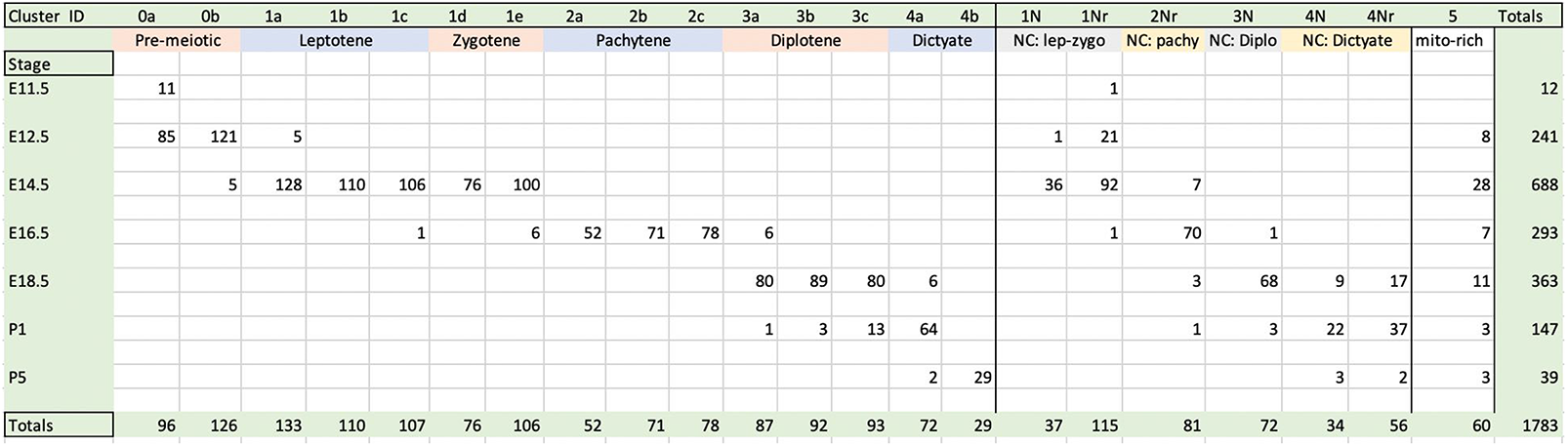
Number of Cells in scRNA clusters vs stage

see Excel File S2.

Excel S4 Genes upregulated in E14.5 and E16.5 nurse cells

Excel File S4

Table S5 “Oocyte” genes upregulated in dictyate oocytes

Excel File S5

### Supplemental Methods

Materials

Table S6

### Supplemental Figures

**Figure S1.**
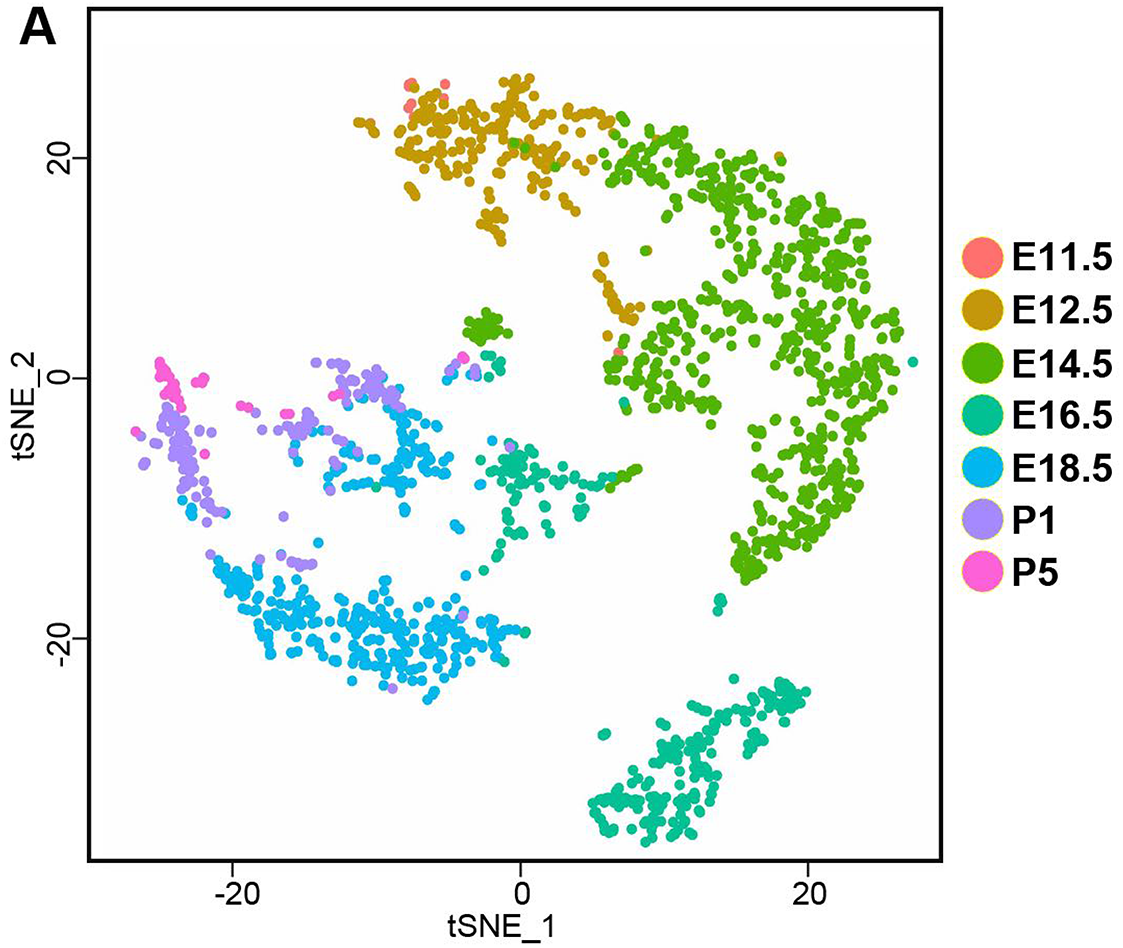
Developmental timing of scRNAseq clusters. tSNE plot identical to that shown in Figure 1C, but color coded to reveal the developmental timing of the component cells (see also Figure 1E and Table S2).

**Figure S2.**
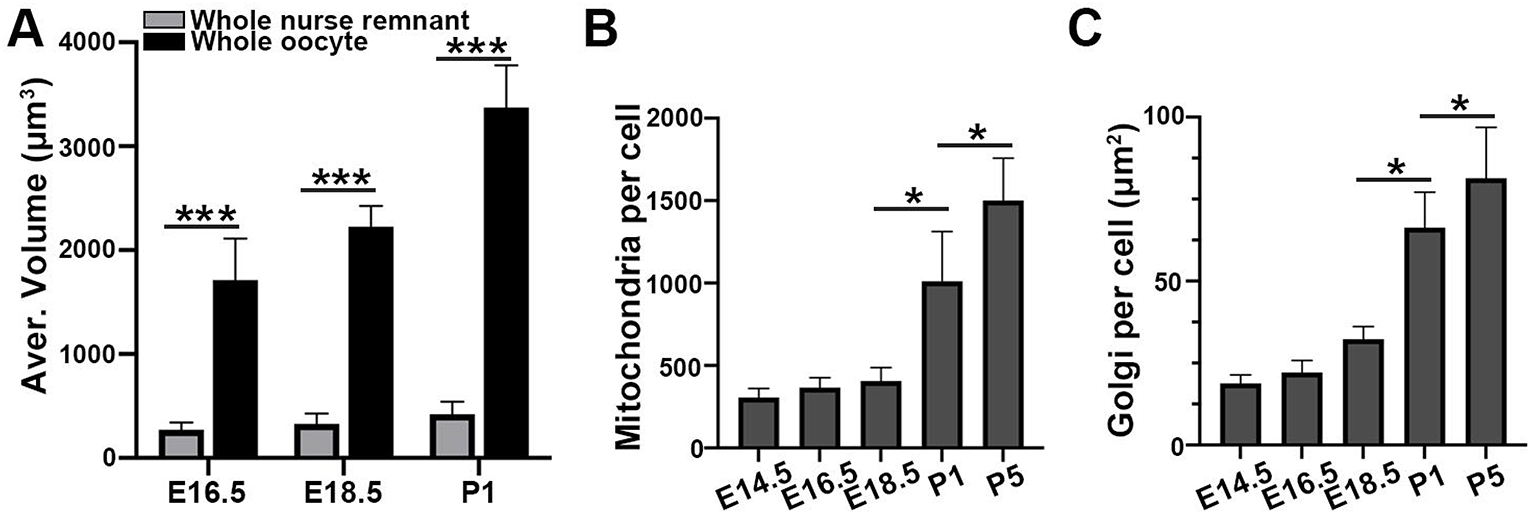
Increase in germ cell volume and organelle content during E14.5-P5. Quantitation of the average volume, mitochondria/cell and Golgi area/cell in the cysts studied in Figure 2.

**Figure S3.**
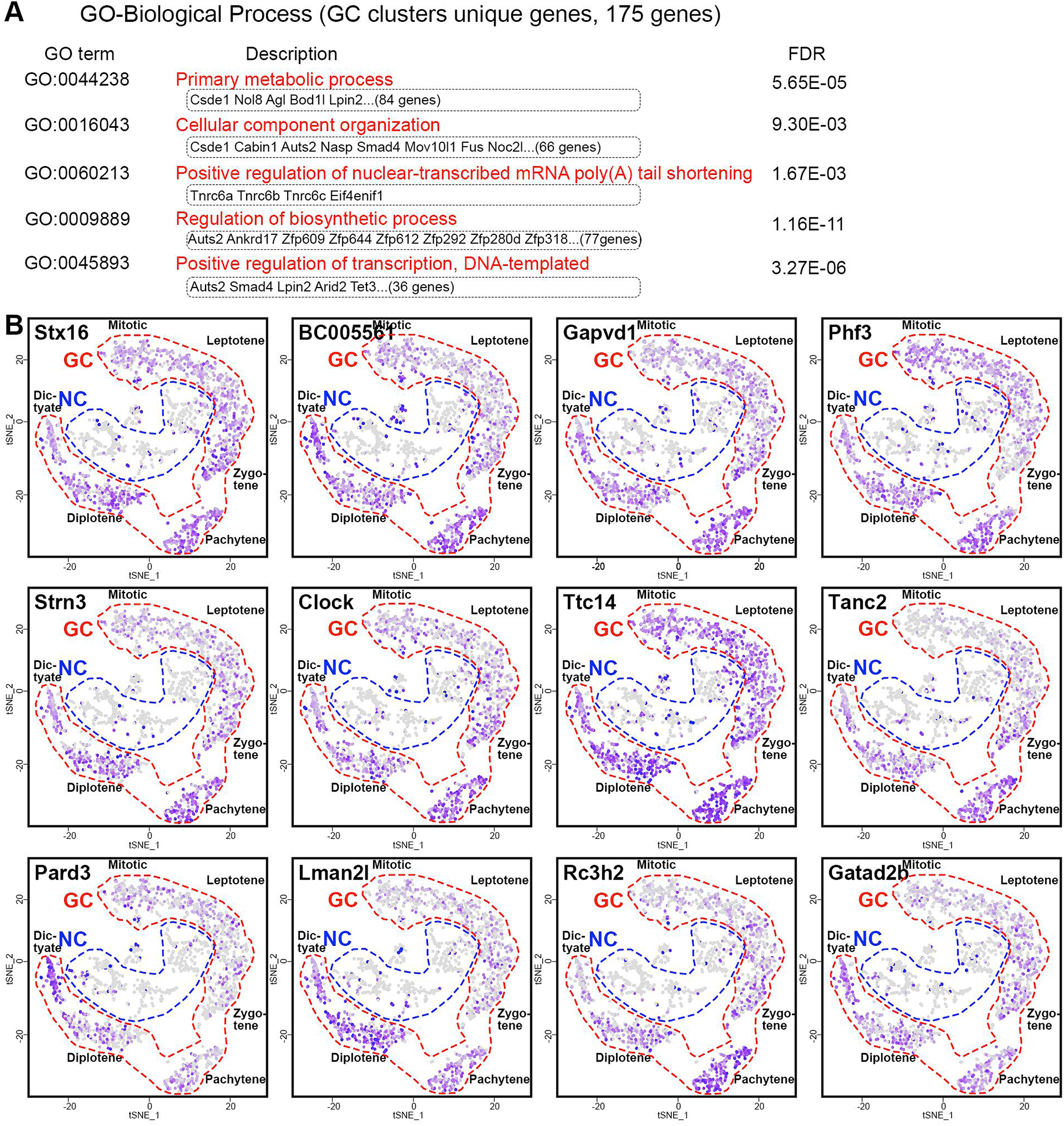
Analysis of genes preferentially expressed in meiotic cells vs nurse cells. A. GO analysis of 175 genes preferentially expressed in full sized germs compared to nurse cells. B. Expression plots of 12 examples of this gene class showing their cell by cell expression on mini tSNE plots. The color indicates level of expression.

**Figure S4.**
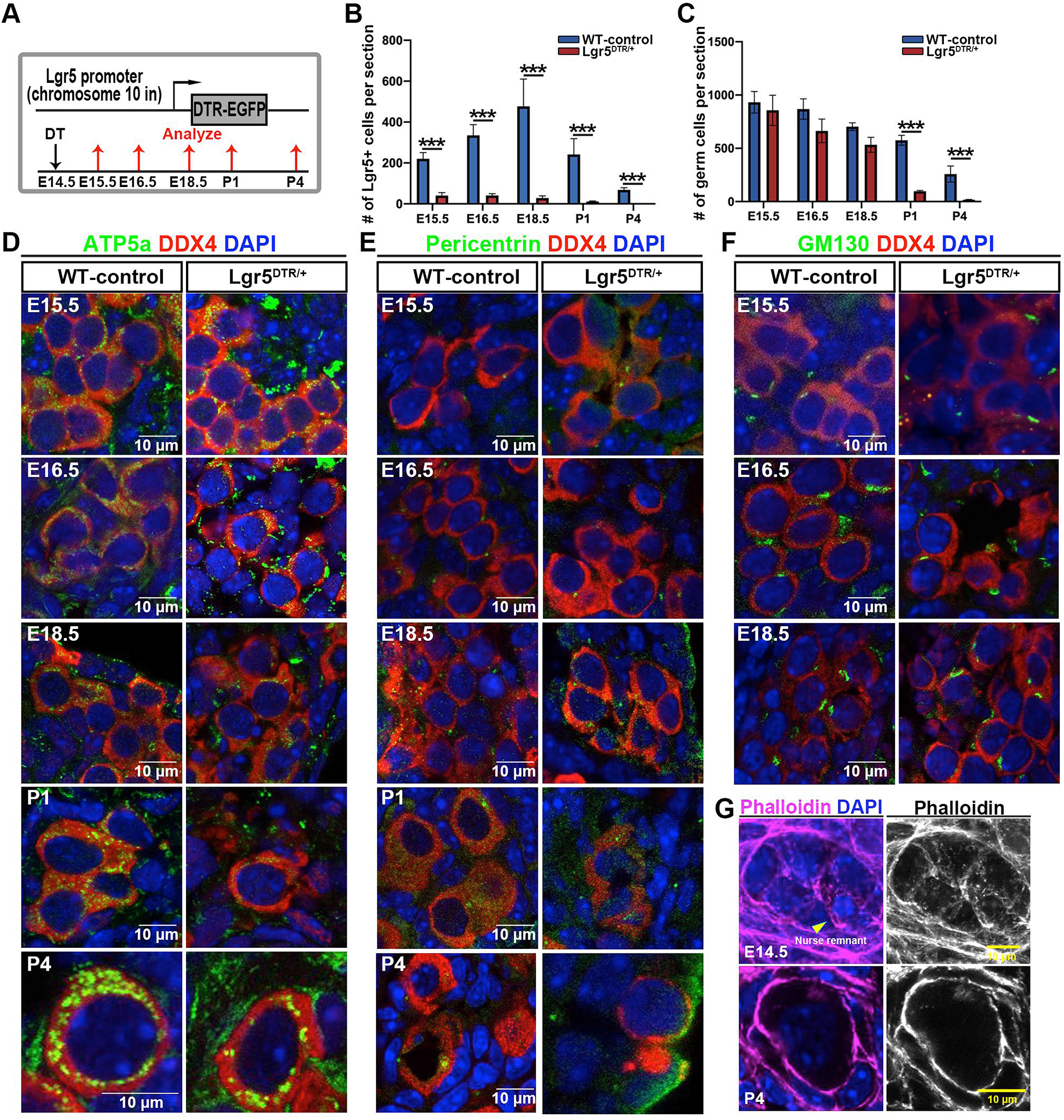
Temporal effects of somatic cell ablation on organelle transfer. A. Scheme for ablation of EPG cells using Lgr5-DTR-EGFP and DT injection. B. Quantitation of Lgr5+ cells in control and EPG ablated ovaries. C. Quantitation of germ cell numbers in control and EPG-ablated cultures. D-F. Examples of germ cells stained to reveal Golgi elements, mitochondria, or centrosomes at E15.5, E16.5, E18.5, P1 and P5. G. E14.5 ovarian cysts (upper) and P4 primordial follicle (lower) stained with phalloidin to reveal actin fibers in germ cells. Aggregations of actin are seen associated with small nurse cell nuclei (nurse remnants, yellow arrowheads). Size bars are indicated.

**Figure S5.**
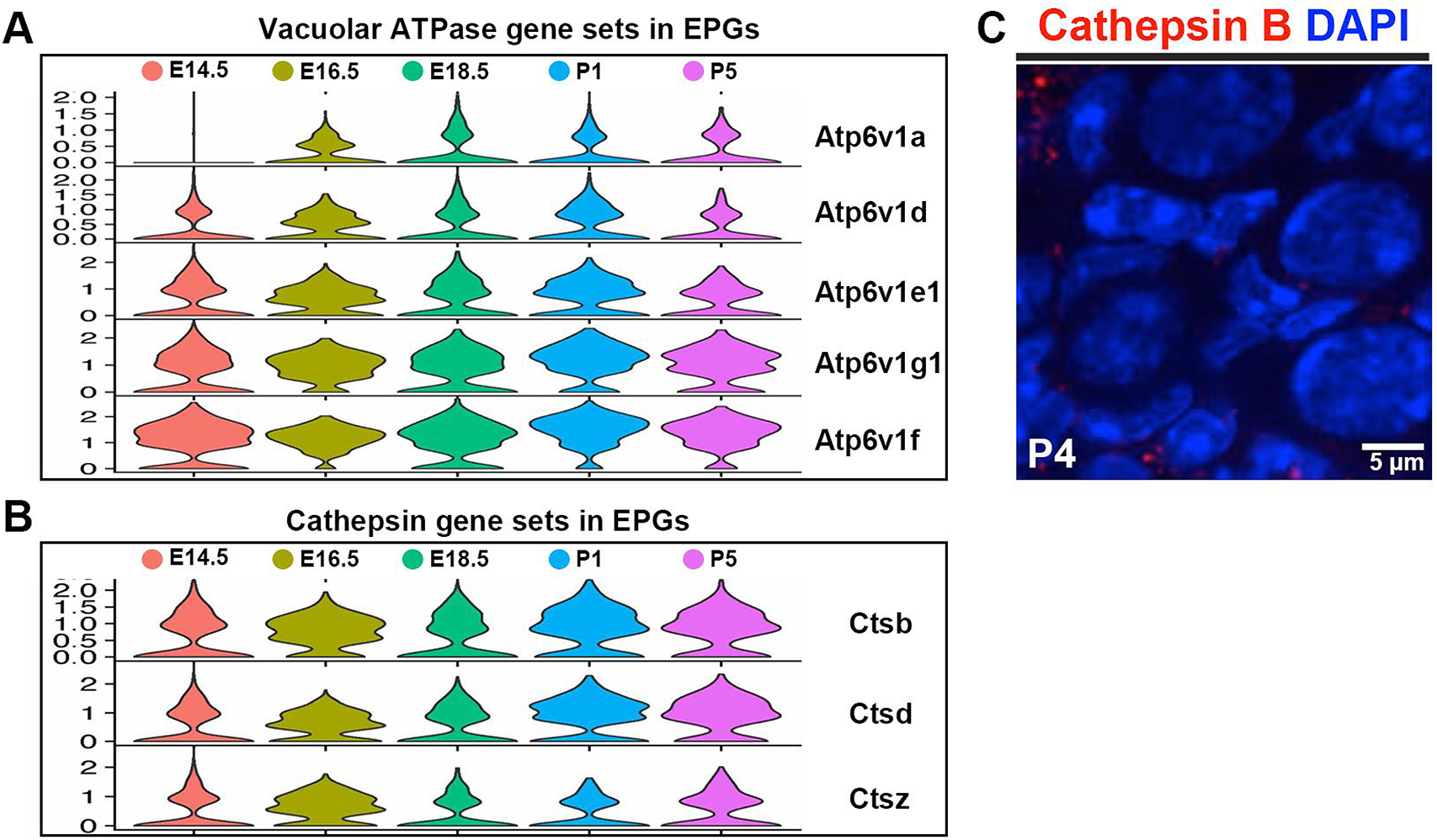
Vacuolar ATPase and Cathepsin gene expression in EPGs. Robust expression of multiple V-ATPase (A) and cathepsin genes (B) in EPGs during E18.5-P1 (data from Niu and Spradling, 2020). C. Cathepsin B expression is no longer detected in P4 ovarian somatic cells.

**Figure S6.**
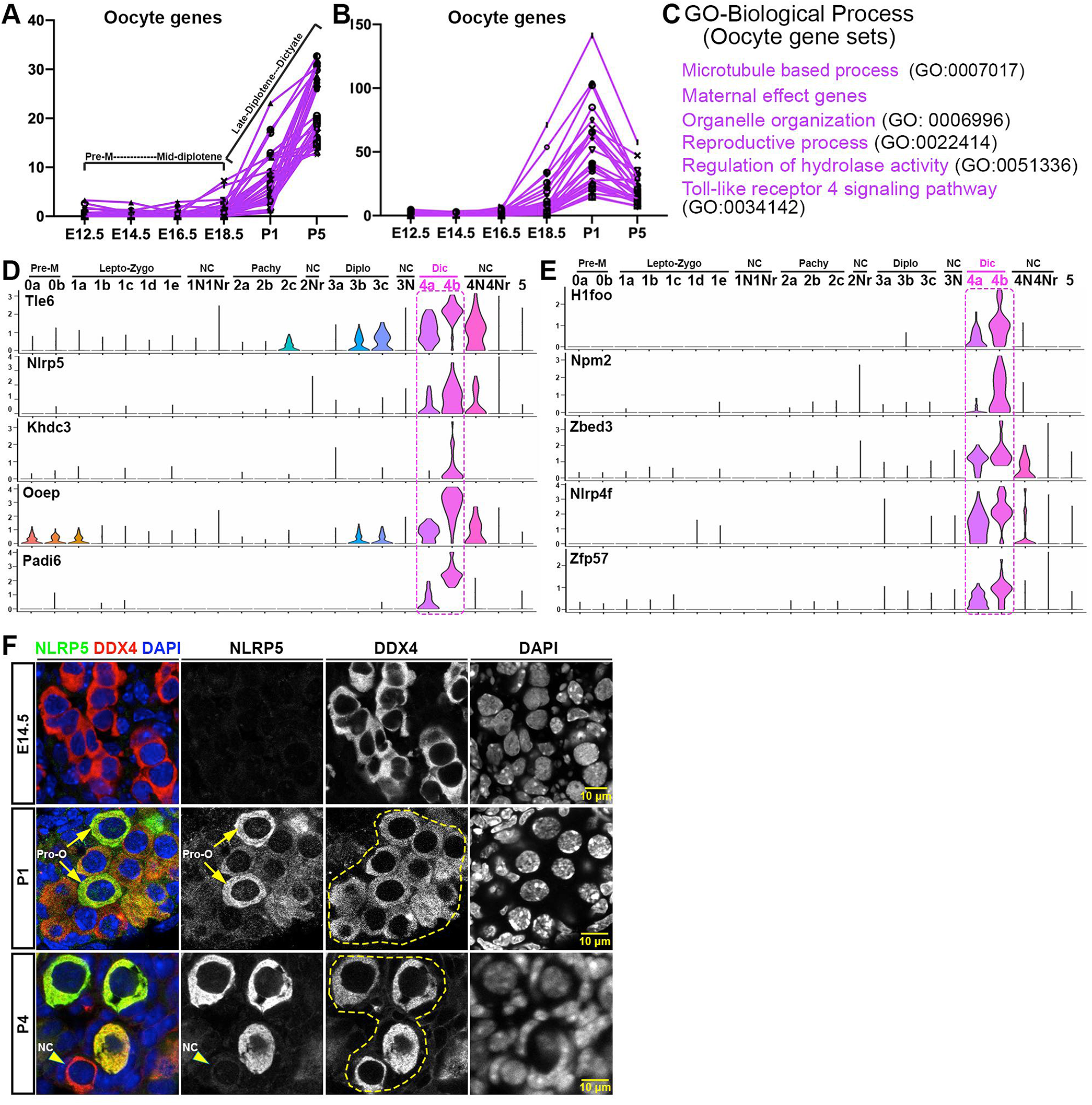
Expression of selected “oocyte” genes. A-B. Expression profiles of selected genes strongly upregulated in late-diplotene and dictyate oocytes that peak at P5 (A) or P1 (B). C. GO analysis of these genes. D-E. Violin plots showing the relative expression of maternal effect genes (a class of “oocyte genes”) in cell clusters. Genes Tle6, Nlrp5, Khdc3, and Ooep make up the subcortical maternal complex (SCMC). (F) Staining of E14.5, P1 and P4 ovarian germ cells with anti-“Mater” (NLRP5) and anti DDX4 antibodies. At P1 and P4, preferential staining of pro-oocytes/oocytes (yellow arrows) and minimal staining of nurse cells (yellow arrowhead) is evident. Size bars are indicated.

**Figure S7.**
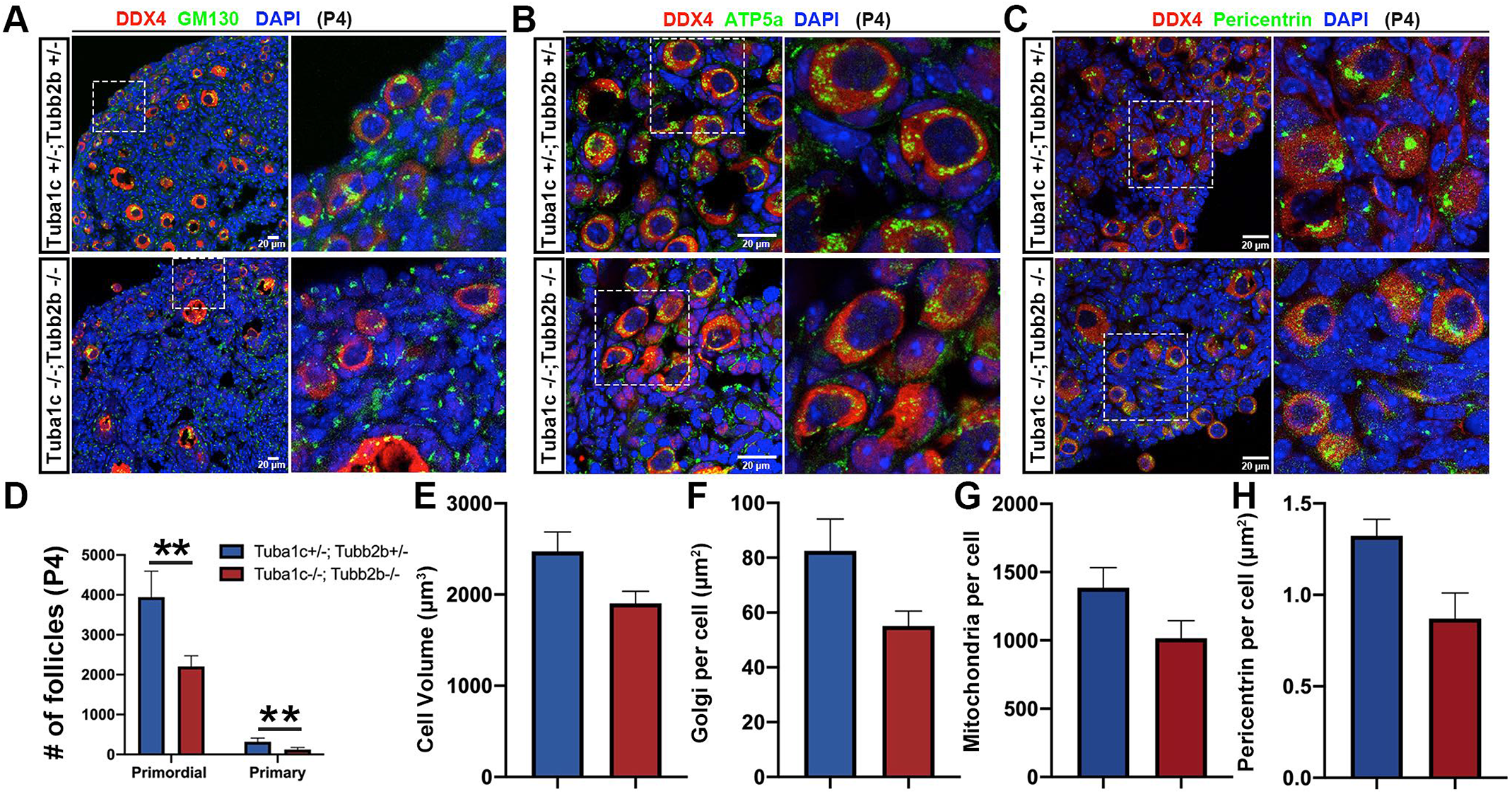
Effects of Tuba1c and Tubb2b knockout on organelle transfer. A-C. Examples of germ cells stained to reveal Golgi elements, mitochondria, or centrosomes at P4 in Tuba1c^+/−^Tubb2b^+/−^ (control) and Tuba1c^−/−^Tubb2b^−/−^ ovaries. D. Quantitation of primordial follicles and primary follicles in P4 ovaries from control and Tuba1c^−/−^Tubb2b^−/−^ animals. E-H. Quantitation of Cell volume (E), Golgi/cell (F), mitochondria/cell (G), Pericentrin (centrosomes)/cell (H) in P4 ovaries from control and Tuba1c^−/−^Tubb2b^−/−^ double animals.

## References

Bittermann E, Abdelhamed Z, Liegel RP, Menke C, Timms A, Beier DR, Stottmann RW. (2019). Differential requirements of tubulin genes in mammalian forebrain development. PLoS Genet. 15: e1008243

Bolivar, J., Huynh, J.R., Lopez-Schier, H., González, C., St Johnston, D., and González-Reyes, A. (2001). Centrosome migration into the Drosophila oocyte is independent of BicD, egl and of the organisation of the microtubule cytoskeleton. Development 128, 1889–1897.

Büning, J. (1994). The Insect Ovary: Ultrastructure, Previtellogenic Growth and Evolution. New York: Chapman & Hall.

de Cuevas, M., Lilly, M. A. and Spradling, A. C. (1997). Germline cyst formation in Drosophila. Annu. Rev. Genet. 31, 405–428.

Cox, R., Spradling, A.C. (2003). A Balbiani body and the fusome mediate mitochondrial inheritance during Drosophila oogenesis. Development 130, 1579–1590.

de Cuevas, M, Spradling, A. C. (1998). Morphogenesis of the Drosophila fusome and its implications for oocyte specification. Development 31, 405–428.

DeLuca, S.Z., Ghildiyal, M., Pang, L. and Spradling, A.C. (2020). Differentiating Drosophila female germ cells initiate Polycomb silencing by regulating PRC2-interacting proteins. eLife 9, e56922

Edson, M.A., Nagaraja, A.K., Matzuk, M.M. (2009). The mammalian ovary from genesis to revelation. Endocr Rev 30, 624–712.

Elkouby, Y. M., Mullins, M. C. (2017). Coordination of cellular differentiation, polarity, mitosis and meiosis—New findings from early vertebrate oogenesis. Dev. Biol., 430, 275–287.

Gondos, B. (1973). Intercellular bridges and mammalian germ cell differentiation. Differentiation 1, 177–182.

Greenbaum, M. P., Yan, W., Wu, M. H., Lin, Y. N., Agno, J. E., Sharma, M., Braun, R. E., Rajkovic, A. and Matzuk, M. M. (2006). TEX14 is essential for intercellular bridges and fertility in male mice. Proc. Natl. Acad. Sci. USA 103, 4982–4987.

Greenbaum, M. P., Iwamori, N., Agno, J. E. and Matzuk, M. M. (2009). Mouse TEX14 is required for embryonic germ cell intercellular bridges but not female fertility. Biol. Reprod. 80, 449–457.

Greenbaum M.P., Iwamori T., Buchold G.M., Matzuk M.M. (2011) Germ cell intercellular bridges. Cold Spring Harb Perspect Biol 3:a005850

Grieder NC, de Cuevas M, Spradling AC. (2000). The fusome organizes the microtubule network during oocyte differentiation in Drosophila. Development. 127: 4253–64.

Grive, K.J., Seymour, K.A., Mehta, R. and Freiman, R.N. (2014) TAF4b promotes mouse primordial follicle assembly and oocyte survival. Dev. Biol., 392, 42–51.

Grive, K.J., Gustafson, E.A., Seymour, K.A., Baddoo, M., Schorl, C., Golnoski, K., Rajkovic, A., Brodsky, A.S. and Freiman, R.N. (2016) TAF4b regulates oocyte-specific genes essential for meiosis. PLos Genet., 12, e1006128.

Haglund K., Nezis I.P., Stenmark H. (2011) Structure and functions of stable intercellular bridges formed by incomplete cytokinesis during development. Commun. Integr. Biol. 4:1–9.

Heasman, J., Quarmby, J.,Wylie, C. C. (1984). The mitochondrial cloud of Xenopus oocytes: The source of germinal granule material. Dev. Biol. 105, 458–469.

Hertig, A. and Adams, E.C. (1967). Studies of the human oocyte and its follicle. J. Cell Biol. 34, 647–675.

Hudson AM, Cooley L. Understanding the function of actin-binding proteins through genetic analysis of Drosophila oogenesis. Annu Rev Genet 2002; 36:455–88.

Huynh, J., St.Johnston, D. (2004). The origin of asymmetry: Early polarization of the Drosophila germline cyst and oocyte. Current Biology 14: R438–49.

Ikami K, Nuzhat N, Lei L. (2017). Organelle transport during mouse oocyte differentiation in germline cysts. Curr Opin Cell Biol. 2017 Feb;44:14–19.

Ikami, K., Nuzhat, N., Abbott, H., Pandoy, R., Haky, L., Spradling, A.C., Tanner H., and Lei, L. (2021). Altered germline cyst formation and oogenesis in Tex14 mutant mice. Biol. Open 10, bio058807.

Jamieson-Lucy, A., Mullins, M. (2019). The vertebrate Balbiani body, germ plasm and oocyte polarity. Curr. Topics in Dev. Biol. 135, 1–34.

Joshi S, Davies H, Sims LP, Levy SE, Dean J. (2007). Ovarian gene expression in the absence of FIGLA, an oocyte-specific transcription factor. BMC Dev Biol 2007; 7:67.

King, R.C. (1970). Ovarian Development in Drosophila melanogaster (Academic Press).

Kloc M, Bilinski S, Dougherty MT, Brey EM, Etkin LD (2004) Formation, architecture and polarity of female germline cyst in Xenopus. Dev Biol 266:43–61.

Koch, E.A. and King, R.C. (1966), The origin and early differentiation of the egg chamber in *Drosophila melanogaster*. J. Morphol. 119, 283–304.

Koch, E. A. and Spitzer, R. H. (1983). Multiple effects of colchicine on oogenesis in Drosophila: induced sterility and switch of potential oocyte to nurse-cell developmental pathway. Cell Tissue Res. 228, 21–32.

Lebo, D.P. V., McCall, K. (2021). Murder on the ovarian express: a tale of non-autonomous cell death in the Drosophila ovary. Cells 10, 1454–73.

Lei, L. and Spradling, A.C. (2013a). Mouse primordial germ cells produce cysts that partially fragment prior to meiosis. Development 140: 2075–2081.

Lei, L. and Spradling, A.C. (2016). Mouse oocytes differentiate through organelle enrichment from sister cyst germ cells. Science 352, 95–99.

Li, L., Baibakov, B., Dean, J. (2008). A subcortical maternal complex essential for preimplantation mouse embryogenesis. Dev. Cell 15, 416–425.

Liang, L.F., Soyal S.M., Dean J. (1997). FIG alpha, a germ cell specific transcription factor involved in the coordinate expression of the zona pellucida genes. Development. 124: 4939–4947.

Lin, Y-T., Capel, B. (2015). Cell fate commitment during mammalian sex determimnation. Curr. Topic in Genet. Dev. 32, 144–52.

Lu K, Jensen L, Lei L, Yamashita YM (2017) Stay connected: a germ cell strategy. Trends Genet 33: 971–978.

Matova, N. and Cooley, L. (2001). Comparative aspects of animal oogenesis. Dev. Biol. 231, 291–320.

Mahowald, A.P., Strassheim, J. M. (1970). Intercellular migration of centrioles in the germarium of Drosophila melanogaster. J. Cell Biol. 45, 306–320.

Mondragon, A.A., Yalonetskaya, A., Ortega, A.J., Zhang, Y., Naranjo, O., Elguero, J., Chung, W.-S., McCall, K. (2019). Lysosomal machinery drives extracellular acidification to direct non-apoptotic cell death. Cell Rpt. 27, 11–19.

Nakamura, S., Kobayashi, K., Nishimura, T., Higashijima, S., and Tanaka, M. (2010). Identification of germline stem cells in the ovary of the teleost medaka. Science 328, 1561–1563.

Nagaoka SI, Nakaki F, Miyauchi H, Nosaka Y, Ohta H, Yabuta Y, Kurimoto K, Hayashi K, Nakamura T, Yamamoto T, Saitou M. (2020). ZGLP1 is a determinant for the oogenic fate in mice. Science 367: eaaw4115.

Niu, W. and Spradling, A.C. (2020). Two distinct pathways of pre-granulosa cell differentiation support follicle formation in the mouse ovary. Proc. Natl. Acad. Sci, USA 117, 20015–20026.

Palozzi, J.M., Swathi, P.J, Hurd, T.R. (2018). Mitochondrial DNA purifying selection in mammals and invertebrates. J. Mol. Biol. 430, 4834–48.

Pepling, M. E. and Spradling, A. C. (1998). Female mouse germ cells form synchronously dividing cysts. Development 125, 3323–3328.

Pepling, M. E. and Spradling, A. C. (2001). Mouse ovarian germ cell cysts undergo programmed breakdown to form primordial follicles. Dev. Biol. 234, 339–351.

Pepling, M., Wilhelm, J.E., O’Hara, A.L., Gephardt, G.W. and Spradling, A.C. (2007). Mouse oocytes within germ cell cysts and primordial follicles contain a Balbiani Body. Proc. Natl. Acad. Sci., 104, 187–192.

Richards, J.S., Pangas, S. (2010). The ovary: basic biology and clinical implications. J. Clin. Invest. 120, 963–72.

Röper K, Brown NH. (2004). A spectraplakin is enriched on the fusome and organizes microtubules during oocyte specification in Drosophila. Curr Biol. 14: 99–110.

Satija R, Farrell JA, Gennert D, Schier AF, Regev A. (2015) Spatial reconstruction of single-cell gene expression data. Nature Biotechnology 33, 495–502.

Stévant I, Kühne F, Greenfield A, Chaboissier MC, Dermitzakis ET, Nef S. (2019). Dissecting Cell Lineage Specification and Sex Fate Determination in Gonadal Somatic Cells Using Single-Cell Transcriptomics. Cell Rep. 26: 3272–3283.

Spradling, A. (1993). Developmental genetics of oogenesis. In The Development of Drosophila melanogaster, M. Bate and A. Martinez-Arias, eds. (Cold Spring Harbor Lab Press), pp. 1–70.

Soh, Y.Q., Junker JP, Gill ME, Mueller JL, van Oudenaarden A, Page DC (2015). A Gene Regulatory Program for Meiotic Prophase in the Fetal Ovary. PLoS Genet. 11, e1005531.

Świątek, P., Urbisz, A.Z. (2017). Architecture and Life History of Female Germ-Line Cysts in Clitellate Annelids. Results and Problems in Cell Differentiation 68, 515–551.

Tharp ME, Malki S, Bortvin A. (2020). Maximizing the ovarian reserve in mice by evading LINE-1 genotoxicity. Nat Commun. 11:330.

Tilly, J.L. (2001). Commuting the death sentence: how oocytes strive to survive. Nat. Rev. Mol. Cell Biol. 2, 838–848.

Wang S, Hassold T, Hunt P, White MA, Zickler D, Kleckner N, Zhang L. 2017. Inefficient crossover maturation underlies elevated aneuploidy in human female meiosis. Cell 168: 977–989 e917

Wang, Z., Liu, C.-Y., Zhao, Y., Dean, J. (2020). FIGLA, LHX8 and SOHLH1 transcription factor networks regulate mouse oocyte growth and differentiation, Nucleic Acids Research 48, 3525–3541.

Wu, D., Dean, J. (2020). Maternal factors regulation preimplantation development in mice. Curr. Top. Dev. Biol. 140, 317–340.

Zheng, G.X. et al. (2017) Massively parallel digital transcriptional profiling of single cells. Nat. Commun. 8, 14049.

